# Seeing speech: neural mechanisms of cued speech perception in prelingually deaf and hearing users

**DOI:** 10.1101/2024.12.06.626971

**Authors:** Annahita Sarré, Laurent Cohen

## Abstract

For many deaf people, lip-reading plays a major role in verbal communication. However, lip movements are by nature ambiguous, so that lip reading does not allow for a full understanding of speech. The resulting language access difficulties may have serious consequences on language, cognitive and social development. Cued speech (CS) was developed to eliminate this ambiguity by complementing lip-reading with hand gestures, giving access to the entire phonological content of speech through the visual modality alone. Despite its proven efficiency for improving linguistic and communicative abilities, the mechanisms of CS perception remain largely unknown. The goal of the present study is to delineate the brain regions involved in cued speech perception and identify their role in visual and language-related processes.

Three matched groups of participants were scanned during two fMRI experiments: Prelingually deaf users of cued speech, hearing users of cued speech, and naive hearing controls. In Experiment 1, we presented videos of silent CS sentences, isolated lip movements, isolated gestures, plus CS sentences with speech sounds and meaningless CS sentences. In Experiment 2, we presented pictures of faces, bodies, written words, tools, and houses in order to understand the contribution of category-specific visual regions to CS perception.

We delineated a number of mostly left-hemisphere brain regions involved in CS perception. We first found that language areas were activated in all groups by both silent CS sentences and isolated lip movements, and by gestures in deaf participants only. Despite overlapping activations when perceiving CS, several findings differentiated experts from novices. As such, CS expertise was associated with a leftward functional lateralization of the lateral occipital cortex, possibly driven by the expert identification of hand gestures. The Visual Word Form Area, which supports the interface between vision and language during reading, did not contribute to CS perception. Moreover, the integration of lip movements and gestures took place in a temporal language-related region in deaf users, and in movement-related regions in hearing users, reflecting their different profile of expertise in CS comprehension and production. Finally, we observed a strong involvement of the Dorsal Attentional Network in hearing users of CS, and identified the neural correlates of the variability in individual proficiency.

Cued speech constitutes a novel pathway for accessing core language processes, halfway between speech perception and reading. The current study provides a delineation of the common and specific brain structures supporting those different modalities of language input, paving the way for further research.

## Introduction

Writing was invented to allow fleeting utterances the potential of enduring over time. In non-logographic writing systems, this is achieved though the visual coding of speech sounds, with some variation in the sound units being written down, which may be phonemes, syllables, etc. Making sounds visible was also prompted by the need to communicate efficiently with deaf people. Unable to perceive speech through the auditory modality, this population heavily relies on lip-reading when encountering spoken language (Desai et al., 2008; Schorr et al., 2005). However, the phonological information carried by the configuration of the mouth are intrinsically ambiguous, such that the words “bark”, “mark” and “park” cannot be distinguished on a visual basis. When no effective strategy is implemented to compensate for this limitation and permit good deciphering of phonological contrasts, the resulting language access difficulties may have serious consequences on language, cognitive and social development (Friedmann and Rusou, 2015; Werker and Hensch, 2015).

In this context, cued speech (CS) was designed by Dr. R. Orin Cornett (Cornett, 1967), with the initial purpose of helping prelingually deaf children improve their reading capacities. Cued speech relies on a set of hand gestures that are designed to counteract the ambiguity of lip-reading, thus giving access to the entire phonological content of speech. As alphabetic scripts are based on a transcription of speech sounds, cued speech improves general linguistic skills of deaf users, notably in reading (Gardiner-Walsh et al., 2020; Trezek, 2017).

In cued speech, words are first decomposed into consonant-vowel (CV) syllables (e.g. “pari” → /pa-ʁi/), sometimes requiring adjustments (e.g. “drakkar” → /d-ʁa-ka-ʁ/ instead of /dʁa-kaʁ/). The identity of each syllable is then conveyed through 3 cues: the lip movements, the position of the hand and the shape of the hand (Figure 1A). The hand assumes one among 8 possible shapes (e.g. index extended and other fingers folded), each shape representing ∼3 consonant phonemes (e.g. /p/, /d/ and /ʒ/). In parallel, the hand is placed in one among 5 possible positions relative to the face (e.g. next to the chin), each position representing ∼3 vowel phonemes (e.g. /a/, /o/, and /œ/). The system is designed such that two syllables sharing the same lip movements will be supplemented by two different hand gestures, allowing an easy differentiation. To a given combination of the three clues thus corresponds a unique syllable, allowing cued speech to convey the utterance’s full phonemic content through the visual modality alone.

**Figure 1.**
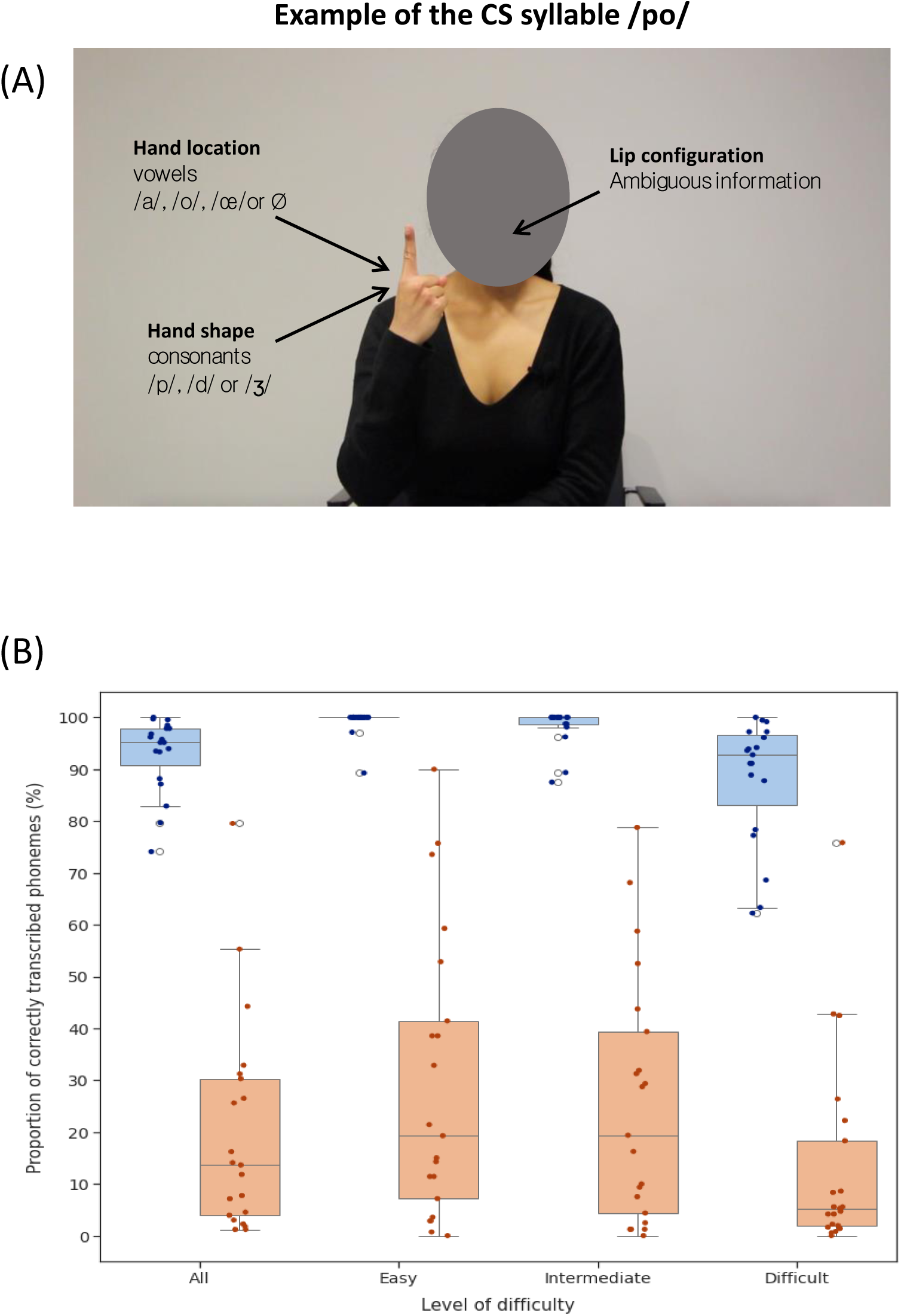
(A) Example syllable from the French cued speech system. CS systems are designed so that syllables sharing the same lip movement are complemented with different hand gestures, allowing for syllabic identification through the visual modality alone. Hand position specify vowels, and hand shape specifies consonants; (B) CS comprehension accuracy, as indexed by the percentage of correctly transcribed phonemes, in deaf (blue) and hearing (orange) CS users, for all sentences and for each level of difficulty. Hearing CS users performed less well, and with larger individual variability, than deaf participants.

The original American CS system was adapted to over 65 languages and dialects (International Academy Supporting Adaptations of Cued Speech (AISAC), 2020), and notably in French where the system is referred to as “Langue française Parlée Complétée” (LfPC). CS is typically used in complement with cochlear implants or, less often, hearing aids.

Naturally CS is not the only way for deaf people to acquire language. The dominant alternative is the use of sign languages, which are fully fledged visual languages distinct from spoken languages and permitting a full linguistic development. When communicating through a given sign language, an ancillary fingerspelling system visually similar to CS can typically be used to convey manually the spelling of words from a spoken language, for instance to communicate proper nouns (Fenlon et al., 2017). Importantly, the use of CS and of a sign language are not incompatible. In fact, studies and guidelines suggest that a bilingual education combining a spoken language supplemented by a CS system and a sign language is beneficial for children (Alegria et al., 1999; Colin et al., 2021).

While sign languages have been the object of a vast number of cognitive neuroscience studies (MacSweeney et al., 2008; Trettenbrein et al., 2021), CS systems have received limited attention from this field. To our knowledge only two studies have been devoted to the brain mechanisms of CS perception, using MRI (Aparicio et al., 2017) and EEG (Caron et al., 2023). Aparicio et al. (2017) presented single words to deaf users of CS in their full CS form, with hand gestures only, and with lip-reading only. The same words were presented to naïve hearing controls in audiovisual form, only auditorily, and with lip-reading only. They observed identical activation of core language areas in both groups, and proposed that the integration of CS gestures and lip-reading cues takes place in the left occipitotemporal junction, particularly in area MT/V5. They also suggested that manual cues may play a larger role than lip-reading in CS perception, in line with indirect behavioral evidence (Attina et al., 2005, 2004; Bayard et al., 2014).

Going beyond those early results, the goal of the present study is to answer fundamental questions on CS perception, using fMRI in three matched groups of participants: prelingually deaf users of cued speech, hearing users of cued speech, and naïve hearing controls. We will explore the following issues and assess the associated predictions.

First, what are the sectors of the visual cortex which are processing hand configurations and lip-reading, and do activation patterns differ between groups, reflecting expertise in CS perception? Concerning hand gestures, the key candidates are the visual word form area (VWFA) and the lateral occipitotemporal cortex (LOTC), two specialized high-level visual areas. Following reading acquisition, the VWFA becomes specialized for the recognition of written letters and words, as shown by a host of neuropsychological and imaging studies (Dehaene et al., 2015). As for the LOTC, it is essential for recognizing all sorts of hand gestures including object manipulation or social communication (Bracci et al., 2018; Wurm and Caramazza, 2019). According to “top-down” theories, the specialization of the VWFA results from its predisposition to interface visual shape analysis with language areas, in which case it should intervene not only in reading but also in CS perception (Price and Devlin, 2011). Conversely, if specialization in the occipitotemporal cortex is mainly driven by the “bottom-up” visual features of stimuli, then the recognition of hand gestures in the context of CS should rely on the LOTC. The predictions are quite different regarding lip-reading. The same mouth configurations are used to convey similar phonological information in both visual and cued speech. Therefore, we predict that the same areas should be involved in lip-reading during CS perception in deaf participants than during visual speech perception in the general population.

Second, how does CS input drive language areas, as compared to the usual visual speech? As CS is only an alternative entry code to the same common language, we predict that the activation of the core, modality independent, language areas (Fedorenko et al., 2024) should be the same in deaf CS users perceiving CS sentences, and in hearing participants perceiving speech. This situation would be similar to the activation of language areas by written and by spoken input (Rueckl et al., 2015). However the two components of CS, when used in isolation, generate distinct predictions. On the one hand, leaving aside possible differences in expertise, pure lip-reading in the absence of gestures and sound, should activate language areas similarly in all groups, as this cue is commonly used for comprehension by both deaf and hearing individuals. On the other hand, perceiving only the gestural component of CS, which is meaningless for naïve participants, may possibly activate language areas exclusively in CS users, again with possible modulations by expertise.

Third, language comprehension often relies on the integration of information conveyed through different acoustic dimensions, like in the integration of speech prosody with syntax (Degano et al., 2024), or through different modalities, like in audiovisual speech perception (Hickok et al., 2018; Ross et al., 2022). Similarly in CS, visual cues from hand position, hand shape, and mouth configuration must be combined to uniquely identify the target syllable. We will try to adjudicate between two options. The convergence may potentially take place either in visual cortices, where each syllable would be represented as a complex gestural combination, or more downstream in language areas, where the different cues would converge to select the appropriate phonological syllable. Both types of convergence of codes have been shown to occur in the field of reading. In alphabetic scripts, upper-case and lower-case letters converge in the occipitotemporal visual cortex (Dehaene et al., 2001), while in Japanese the logographic kanji and the syllabic kana scripts converge in left temporal language areas (Nakamura et al., 2005).

Fourth, we will address the potentially complex case of the hearing CS users. These participants typically master CS production better than perception, as they use CS to address deaf relatives or people whom they assist, but are rarely addressed to in silent CS. They also generally learn CS as adults. We will therefore try to clarify where this group stands with respect to the others two, and whether hearing CS users show important individual variability.

To address these questions, we scanned deaf CS users, hearing CS users and hearing CS naïve controls during two fMRI experiments. In Experiment 1, we presented videos of different types of sentences in full or degraded French CS. In Experiment 2, we presented pictures of faces, bodies, written words, tools, and houses in order both to understand the contribution of category-specific visual regions to CS perception, and to identify potential modification of the functional layout of ventral occipitotemporal cortex (VOTC) in CS users. MRI acquisition was preceded by a questionnaire on deafness and language history, and by a pretest evaluating the mastery of CS of deaf and hearing CS users.

## Methods

### Participants

We recruited 60 healthy volunteers: 19 prelingually and severely/profoundly deaf CS users, 21 hearing CS users and 20 hearing controls with no knowledge of CS. Participants were 18-65 years old French native speakers. All participants were right-handers according to the Edinburgh inventory (Oldfield, 1971) except for two left-handed deaf participants and one ambidexter hearing user. They had no history of neurological or psychiatric disorders, and all had normal or corrected to normal vision. They did not present contra-indications to MRI, which constituted a particular challenge as deaf CS users typically carry an MRI-incompatible cochlear implant. Only participants with no or removable hearing aids were recruited. For the two groups of CS users, CS comprehension and production were both assessed before the MRI session. For the reason discussed above, deaf participants were required to have a good level in CS comprehension, and hearing participants to master CS production. Recruitment of CS users was done through social networks and mailing lists of CS associations.

Before the MRI session, the participants were asked to fill an initial questionnaire at home containing general demographic questions for all three groups of participants. Deaf CS users were asked additional questions focusing on the etiology and severity of deafness, history of language acquisition, hearing aids and on the daily use of CS. Hearing CS users answered questions on the learning and use of CS in daily life.

Deaf participants were severely (n=2), profoundly (n=16) or totally (n=1) deaf. All were pre-linguistically deaf, including 16 out of 19 congenital deafness. All reported standard ages of reading acquisition (5.15±1.01 years old), with average to good current reading capacities. Hearing users and controls had normal hearing, except for one hearing CS user who had moderate deafness. The three groups were matched in age (Deaf: 35.26±8.12 yo; hearing: 35.52±11.31 yo; controls: 34.35±10.72 yo) and education level (university degree: deaf: n=17; hearing: n=19; controls: n=18). The sex ratio was similar in the deaf and control groups (Deaf: 12 F; controls: 13 F), while the hearing users of CS included slightly more females (16 F). CS users differed in their age of CS learning (Deaf: 2.84±1.71 yo; hearing: 24.95±10.49 yo). All but two users practiced CS at least once a month, with more hearing participants practicing daily or several times a week (Deaf: n=11; hearing: n=16). The two remaining participants were proficient deaf early CS users and scored high on the pretest. Moreover, 15 deaf and 13 hearing users declared having knowledge on French Sign Language, with a later age of learning than for CS (Deaf: 15.67±9.24 yo; hearing: 24.61±10.98 yo) and varying self-declared levels in comprehension and production but a rather frequent use by deaf participants (12 using French Sign Language at least once a month).

All information was provided in written form identically to the 3 groups of participants. The research was approved by the institutional review board “Comité de Protection des Personnes” Est-III (N° CPP 20.11.05). All participants provided informed written consent in accordance with the Declaration of Helsinki.

### Behavioral assessment

The proficiency of CS users in CS comprehension and production was assessed before the MRI session. Comprehension was tested through a short dictation test, where the participants had to write down 12 CS sentences presented as silent videos. Each sentence was presented twice, and participants had to respond after each viewing. CS production was tested by asking participants to transpose 12 written sentences into CS. The accuracy and fluency of responses were checked by the experimenter. For CS comprehension, responses were assessed by computing for each sentence the percentage of correctly transcribed phonemes. Spelling mistakes were disregarded as long as they transcribed the correct phoneme. Results were then averaged across sentences.

For both tests, sentences were distributed into three levels of difficulty. “Easy” sentences were short, used the present tense, included only frequent and semantically predictable words, and CV syllables. “Intermediate” sentences were short, used various tenses, and included frequent but less predictable words and one complex syllabic pattern (V-CCV, V-CVC or VC-CV) (Alegria et al., 1999). “Difficult” sentences were longer, used various tenses, included less frequent and less predictable words, and one complex syllabic pattern.

### Image acquisition and preprocessing

MRI data were acquired on a Siemens 3T Prisma system at the CENIR imaging center (Paris Brain Institute), using 20 (Experiment 1) and 64 (Experiment 2) channels head coils. To record fMRI data, we used the multi-echo multi-band approach to have high SNR and coverage of the areas sensitive to signal dropout, particularly the lower part of the temporal lobes, while keeping good spatiotemporal resolution. Sequence parameters were TR/TEs/FA = 1660ms/14.2ms, 35.39ms, 56.58ms/74°, isotropic voxel size of 2.5mm, 60 slices, acceleration factors were multi-band=3 and iPat(GRAPPA)=2. In most participants, pulse oximeter and respiration belt signals were recorded and used for denoising of the data. The anatomical image was a 3DT1 with 1mm isotropic voxels, using an MPRAGE sequence.

The anatomical image was segmented and normalized to the MNI space using CAT12 (Gaser, 2020). Minimal preprocessing was then conducted with AFNI library (Cox, 1996; Cox and Hyde, 1997) using afni_proc.py wrapper to perform temporal despiking (despike), slice timing correction (tshift), and movement correction (volreg). The volume registration was computed on the first (shortest) echo, and applied to all echos, where the target for the registration is the MIN_OUTLIER volume, corresponding the volume with minimal movement. The 3 echos were optimally combined with TEDANA library (The tedana Community et al., 2021). First, a T2* map was computed using all echos, then the echos were combined with a weighted sum where the weights were a combination of TE and T2*. All subsequent steps used SPM12. Optimally combined volumes were coregistered to the anatomical scan, and normalized to MNI space using the deformation field computed for the anatomical scan. Finally, data were smoothed using a Gaussian kernel of 4 mm FWHM.

A set of noise regressors were derived from the preprocessing using TAPAS/PhysIO (Kasper et al., 2017), derived from cardiac and respiratory recordings (RETROICOR), cardiac recordings (HRV), respiratory recordings (RVT), white-matter and CSF time series, and PCA to reduce their dimensionality (Noise ROI), realignment parameters, their derivatives, and squared parameters and derivatives, and stick regressors to scrub data from volumes with FD>0.5mm.

### Statistical analyses

For single-subject analyses, General Linear Models (GLMs) were created and estimated for each experiment and participant using SPM12, with a regressor for each experimental condition, plus regressors for targets and motor responses for Experiment 2, as well as movement and physiological regressors. For group-level analyses, individual contrast images were entered in t-test models for the different comparisons of interest. Unless stated otherwise, the statistical threshold was set to p<0.001 voxel-wise, and q<0.05 cluster-wise FDR-corrected for multiple comparisons across the whole brain. Cerebellar activations are only reported in the tables.

Some analyses of the data of Experiment 1 were carried out in individual VOTC regions of interest (ROI) defined on the basis of Experiment 2. To define those ROIs, we first identified the peak coordinate of the following group-level contrasts : the conjunctions of activations in all three groups of Faces > Other categories to identify the Fusiform Face Area (FFA), of Bodies > Other categories to identify the Extrastriate Body Area, and of Words > Other categories to identify the VWFA. For each contrast, we selected a peak in each hemisphere, except for the strongly left-lateralized VWFA, for which we selected the symmetric right-hemisphere coordinates. Around each peak, we defined a sphere of 8 mm radius. Finally, within each of those spheres, we identified the individual local peak activation in the corresponding contrast. Individual 4 mm radius spheres centered on those peaks were used as ROIs in which data from Experiment 1 were sampled.

### Experiment 1: Brain activation during cued-speech perception

The experiment included 5 conditions : (1) Sentences in full CS (with sound), (2) Sentences in full CS (silent), (3) Sentences in lip-reading only (silent), (4) Sentences with only the gestural part of CS (silent), (5) Meaningless pseudo-sentences in full CS (silent), as well as baseline periods with a fixation cross.

For each condition, we created 16 different stimuli. Each of these sentence or pseudo-sentence included 19 CS gestures, and lasted about 5 seconds. Pseudo-sentences were derived from a subset of the real sentences from conditions 1-4, in which open-class words were modified into pseudowords. Each trial started with a 0.2 s image of the background without the coder, followed by a 0.2 s smooth transition, then 2 stimuli from the same condition, separated by a 0.2 s transition. The trial ended with a 0.2 s transition and a 0.2 s background image, for a total duration of 13 s per trial. Each stimulus was used once, resulting in 8 trials per condition.

Moreover, 8 baseline periods of 13 s each were displayed. They consisted of a black fixation cross at the location of the coder’s chin, on an image of the background.

The 40 trials (8 for each of the 5 conditions) and 8 baseline periods were combined in two pseudo-random orders, such that no condition could be presented more than twice in a row, and that two baseline periods never occurred consecutively. Half the subjects were randomly presented with each order. The experiment had a total duration of about 11 minutes, and began and ended with additional baseline periods.

Material for the pretest and the experiment were recorded at the Paris Brain Institute with a professional CS coder and edited using iMovie. Stimuli presentation was done using Psychtoolbox Version 3 (Brainard, 1997) in MATLAB R2019b.

Participants were asked to pay attention to the stimuli and to understand them as much as they could.

### Experiment 2: Brain activation during visual objects perception

In Experiment 2, static images were presented to participants, distributed among 5 visual object categories, each represented by 20 pictures: faces, bodies, French words, houses and tools (for a full description of stimuli see Zhan et al., 2023).

The experiment consisted of 12 blocks for each category of stimuli, arranged in a random order. In each block, all 20 pictures from the considered category were presented, each time in a different random order. Each picture was presented for 100 ms, followed by a 200 ms blank screen with a fixation cross. Blocks had thus a duration of 6 s, and were separated with 6 s of fixation baseline, for a total session duration of 12 minutes.

Stimuli presentation was done using Psychtoolbox Version 3 (Brainard, 1997) in MATLAB R2019b.

A white fixation cross was present at the center of the screen throughout the experiment. Participants were asked to fixate the cross, to detect the picture of a star which replaced a picture in half the blocks of each category, and to press a response button as rapidly and as accurately as possible.

## Results

### Behavioral assessment of cued speech comprehension

The deaf and hearing CS users were presented with silent CS sentences and asked to write them down. Each sentence was presented twice, and two transcriptions were required.

Averaging both answers, Deaf CS users responded accurately (in each sentence, 92.91±7.27% of phonemes were correctly transcribed), showing little variation across subjects (Figure 1B). The performance of hearing CS users was on average much lower (19.72±20.56%; Mann-Whitney U=398, p<0.005), and more heterogeneous (Levene’s test for variance comparison: F=7.07, p<0.05). The superiority of Deaf over hearing CS users prevailed separately for the 3 levels of sentence difficulty (easy: U=398, p<0.005; medium: U=399, p<0.005; hard: U=396, p<0.005).

Neither group showed a significant difference between the “Easy” and the “Intermediate” conditions (p>0.05), while comparing “Intermediate” and “Difficult” conditions showed a significant effect in the deaf (U=314.5, p<0.005) but not in the hearing group. Both groups showed a significant difference between the “Easy” and “Difficult” conditions (Deaf: U=335.5, p<0.005; hearing: U=301.5, p<0.05).

Looking for differences between the first and second responses, we observed that, in the deaf group, the second attempt was significantly better than the first, for all sentences together (U=90.5, p<0.01), and for hard sentences separately (U=96, p<0.05). No difference between attempts was found in the hearing group.

### Experiment 1: Cued speech perception

In order to identify regions activated during the perception of CS, we presented silent CS sentences (henceforth called the “Sentences” condition), silent sentences with only lip-reading cues (“Lip-reading” condition), silent sentences with only gestural cues (“Gestures” condition), CS sentences presented with the corresponding speech sound (“Audible sentences” condition), and silent CS sentences made up of meaningless words (“Pseudo-sentences” condition).

#### Activation by silent sentences in cued speech

In order to delineate the overall set of regions activated during CS perception, we first compared Sentences > baseline (Figure 2; Table 1).

**Figure 2.**
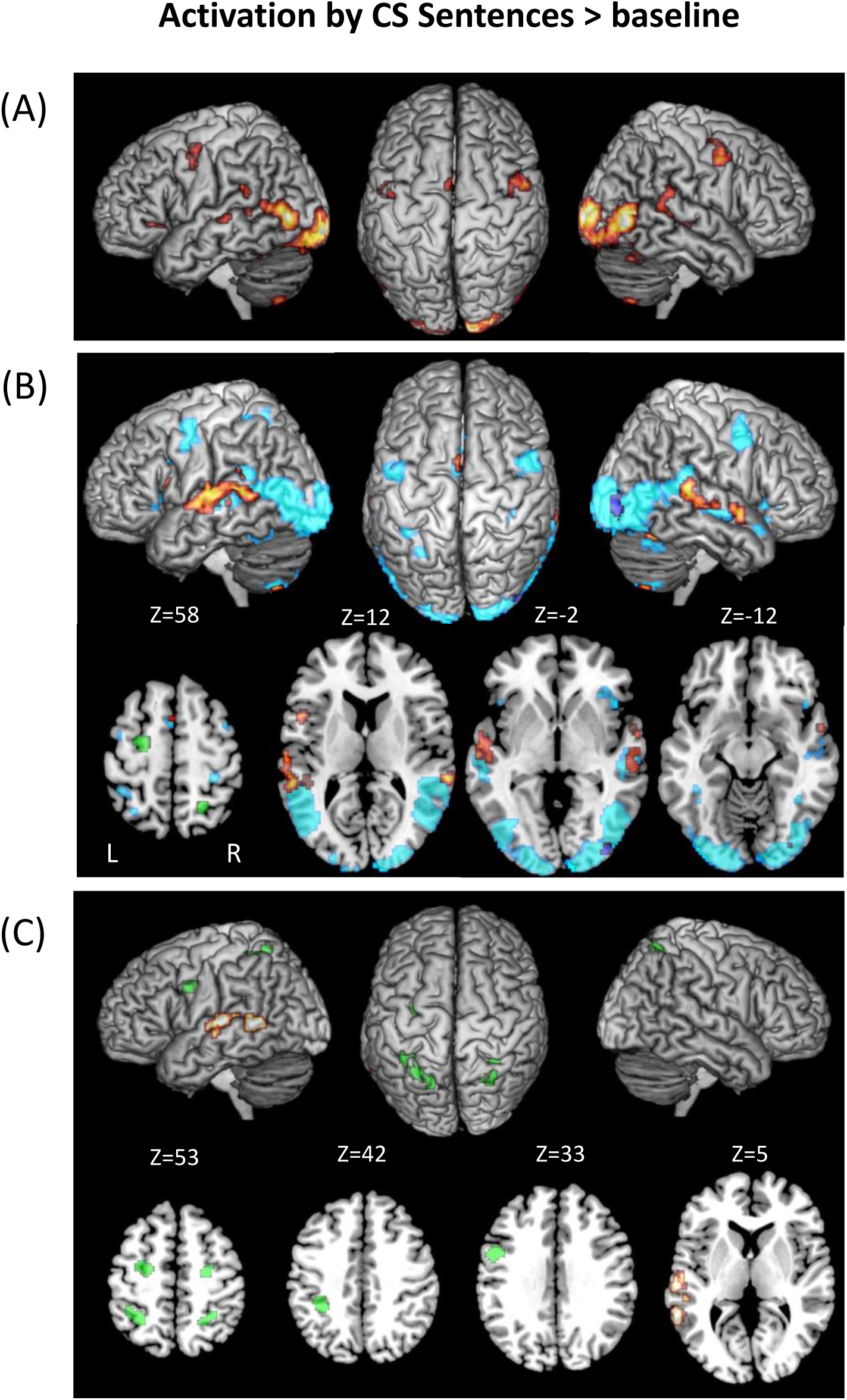
Activation by LPC Sentences > baseline. (A) Activations common to the three groups; (B) Activations in Controls (light blue), in Controls > Deaf (blue), in Deaf > Controls (hot) and in Hearing > Controls (green); (C) Activations in Deaf > Hearing (hot) and Hearing > Deaf (green).

**Table 1.** Sentences > baseline. Hemisphere and anatomical regions, MNI coordinates and Z score of peak activations. Voxelwise threshold p<0.001, clusterwise threshold p<0.05 FDR-corrected over the whole brain. The contrast Controls > Hearing (masked by Controls) showed no significant activation. L=left; R=right; a=anterior; p=posterior; LOTC=lateral occipitotemporal cortex; STS=superior temporal sulcus; STG=superior temporal gyrus; IFG=inferior frontal gyrus; SMA=supplementary motor area; IPS=intraparietal sulcus; FEF=frontal eye field

| Region | Conjunction 3 groups |  |  |  | Deaf > Controls |  |  |  | Deaf > Hearing |  |  |  | Hearing > Deaf |  |  |  | Controls > Deaf |  |  |  | Hearing > Controls |  |  |  |
| --- | --- | --- | --- | --- | --- | --- | --- | --- | --- | --- | --- | --- | --- | --- | --- | --- | --- | --- | --- | --- | --- | --- | --- | --- |
|  | x | y | z | Z | x | y | z | Z | x | y | z | Z | x | y | z | Z | x | y | z | Z | x | y | z | Z |
| L Calcarine sulcus | -12 | -101 | -7 | 7.66 |  |  |  |  |  |  |  |  |  |  |  |  |  |  |  |  |  |  |  |  |
| R Calcarine sulcus | 12 | -101 | 3 | > 8 |  |  |  |  |  |  |  |  |  |  |  |  |  |  |  |  |  |  |  |  |
| L Inferior occipital gyrus | -20 | -96 | -7 | 7.23 |  |  |  |  |  |  |  |  |  |  |  |  |  |  |  |  |  |  |  |  |
| R Inferior occipital gyrus | 22 | -96 | 3 | 6.99 |  |  |  |  |  |  |  |  |  |  |  |  | 40 | -86 | -4 | 4.15 |  |  |  |  |
| L LOTC | -48 | -71 | 3 | 6.83 |  |  |  |  |  |  |  |  |  |  |  |  |  |  |  |  |  |  |  |  |
| R LOTC | 48 | -68 | -2 | > 8 |  |  |  |  |  |  |  |  |  |  |  |  |  |  |  |  |  |  |  |  |
| L Fusiform gyrus | -42 | -44 | -20 | 5.78 |  |  |  |  |  |  |  |  |  |  |  |  |  |  |  |  |  |  |  |  |
| R Fusiform gyrus | 40 | -48 | -14 | 4.66 |  |  |  |  |  |  |  |  |  |  |  |  |  |  |  |  |  |  |  |  |
| L aSTS/STG | -65 | -26 | 3 | 3.62 |  |  |  |  | -60 | -11 | 3 | 3.39 |  |  |  |  |  |  |  |  |  |  |  |  |
| R aSTS/STG | 52 | -16 | -7 | 3.9 | 58 | 2 | -7 | 4.28 |  |  |  |  |  |  |  |  |  |  |  |  |  |  |  |  |
| L pSTS/STG | -48 | -46 | 10 | 5.03 | -58 | -41 | 10 | 5.09 | -52 | -34 | 8 | 4.46 |  |  |  |  |  |  |  |  |  |  |  |  |
| R pSTS/STG | 50 | -36 | 6 | 5.75 | 65 | -36 | 8 | 4.84 |  |  |  |  |  |  |  |  |  |  |  |  |  |  |  |  |
| L Middle temporal gyrus |  |  |  |  |  |  |  |  | -62 | -48 | 6 | 4.17 |  |  |  |  |  |  |  |  |  |  |  |  |
| L Supramarginal gyrus | -45 | -38 | 26 | 3.87 |  |  |  |  |  |  |  |  |  |  |  |  |  |  |  |  |  |  |  |  |
| L IFG (pars opercularis) |  |  |  |  | -50 | 12 | 13 | 4.16 |  |  |  |  |  |  |  |  |  |  |  |  |  |  |  |  |
| L IFG (pars orbitalis) | -38 | 29 | -2 | 3.53 |  |  |  |  |  |  |  |  |  |  |  |  |  |  |  |  |  |  |  |  |
| L Precentral gyrus | -42 | -4 | 50 | 4 |  |  |  |  |  |  |  |  | -50 | 4 | 30 | 4.63 |  |  |  |  |  |  |  |  |
| R Precentral gyrus | 50 | 2 | 46 | 4.27 |  |  |  |  |  |  |  |  |  |  |  |  |  |  |  |  |  |  |  |  |
| L/R SMA | -2 | 6 | 56 | 3.72 | -2 | 4 | 68 | 3.76 |  |  |  |  |  |  |  |  |  |  |  |  |  |  |  |  |
| L Inferior parietal lobule |  |  |  |  |  |  |  |  |  |  |  |  | -35 | -38 | 43 | 5.48 |  |  |  |  |  |  |  |  |
| L IPS |  |  |  |  |  |  |  |  |  |  |  |  | -20 | -58 | 58 | 4.87 |  |  |  |  |  |  |  |  |
| R IPS |  |  |  |  |  |  |  |  |  |  |  |  | 32 | -46 | 48 | 4.73 |  |  |  |  | 20 | -58 | 63 | 3.6 |
| L FEF |  |  |  |  |  |  |  |  |  |  |  |  | -28 | -6 | 56 | 5.39 |  |  |  |  | -22 | -11 | 56 | 4.88 |
| R FEF |  |  |  |  |  |  |  |  |  |  |  |  | 28 | -6 | 46 | 4.76 |  |  |  |  |  |  |  |  |
| L Cerebellum | -10 | -74 | -44 | 4.05 |  |  |  |  |  |  |  |  |  |  |  |  |  |  |  |  |  |  |  |  |
|  | -30 | -66 | -54 | 3.54 |  |  |  |  |  |  |  |  |  |  |  |  |  |  |  |  |  |  |  |  |
| R Cerebellum | 30 | -64 | -57 | 4.4 | 28 | -66 | -57 | 4.37 |  |  |  |  |  |  |  |  |  |  |  |  |  |  |  |  |
|  | 12 | -76 | -44 | 4.21 | 28 | -66 | -57 | 4.37 |  |  |  |  |  |  |  |  |  |  |  |  |  |  |  |  |
|  | 42 | -58 | -30 | 4.47 | 35 | -61 | -24 | 5.37 |  |  |  |  |  |  |  |  |  |  |  |  |  |  |  |  |
|  | 28 | -61 | -27 | 3.86 |  |  |  |  |  |  |  |  |  |  |  |  |  |  |  |  |  |  |  |  |

The conjunction of this contrast across the three groups showed common activation (Figure 2A), (1) in bilateral visual regions, including the occipital poles, lateral and inferior occipital cortex, and fusiform gyri; (2) in language-related areas, including the left-hemispheric inferior frontal gyrus (IFG) left supramarginal gyrus (SMG), and supplementary motor area (SMA); and the bilateral superior temporal sulcus (STS), posterior superior temporal gyrus (pSTG), precentral gyrus, and cerebellum.

We then compared CS users to controls (Figure 2B). First, the comparison of Deaf participants > Controls (masked by Deaf > baseline), showed activation in the left IFG, the left predominant STS/STG and the SMA. The opposite contrast of Controls > Deaf (masked by Controls > baseline) only showed right inferior occipital activation. Second, the contrast of Hearing > Controls (masked by Hearing > baseline) activated the left frontal eye field (FEF) and the right intraparietal sulcus (IPS). The opposite contrast of Controls > Hearing (masked by Controls > baseline) showed no significant activation.

We then compared the two groups of CS users. The contrast of Deaf > Hearing (masked by Deaf > baseline) activated the left STG and middle temporal gyrus (MTG), plus the right pSTS just below the threshold for cluster extent (26 voxels). The opposite comparison of Hearing > Deaf (masked by Hearing > baseline), showed activation in the bilateral FEF and IPS regions already partially present in the Hearing > Controls comparison, in the left precentral gyrus, plus the right inferior occipital cortex at a lower voxelwise threshold (p<0.01) (Figure 2C).

In summary, we found activation in vision- and language-related areas common to all groups, plus three differences among groups. First, there was stronger activation in users of CS as compared to controls, in frontal and temporal language areas in the deaf group, and in temporal regions only in the hearing group. Second, there was stronger activation in a bilateral frontoparietal IPS/FEF network in the hearing than in both the deaf and the control groups. Third, activation was stronger in both the hearing and control groups than in deaf participants in the right inferior occipital cortex, the only difference among groups observed in the visual cortex.

The fact that controls activated language areas, albeit more weakly than deaf participants, may come as a surprise. In the absence of any comprehension of the gestural code by controls, lip-reading appeared as the likely explanation, which we assessed in the following analyses.

#### Activation by Lip-reading and by Gestures

We then contrasted conditions with only partial CS information (Lip-reading or Gestures) minus baseline (Figure 3; Suppl tables 1-2), in order to determine (i) whether such degraded information was sufficient to activate language areas and the IPS/FEF attention-related network, (ii) whether distinct parts of the visual cortex were preferentially involved in the processing of lip-reading and gestures, and (iii) how groups differed in those respects.

**Figure 3.**
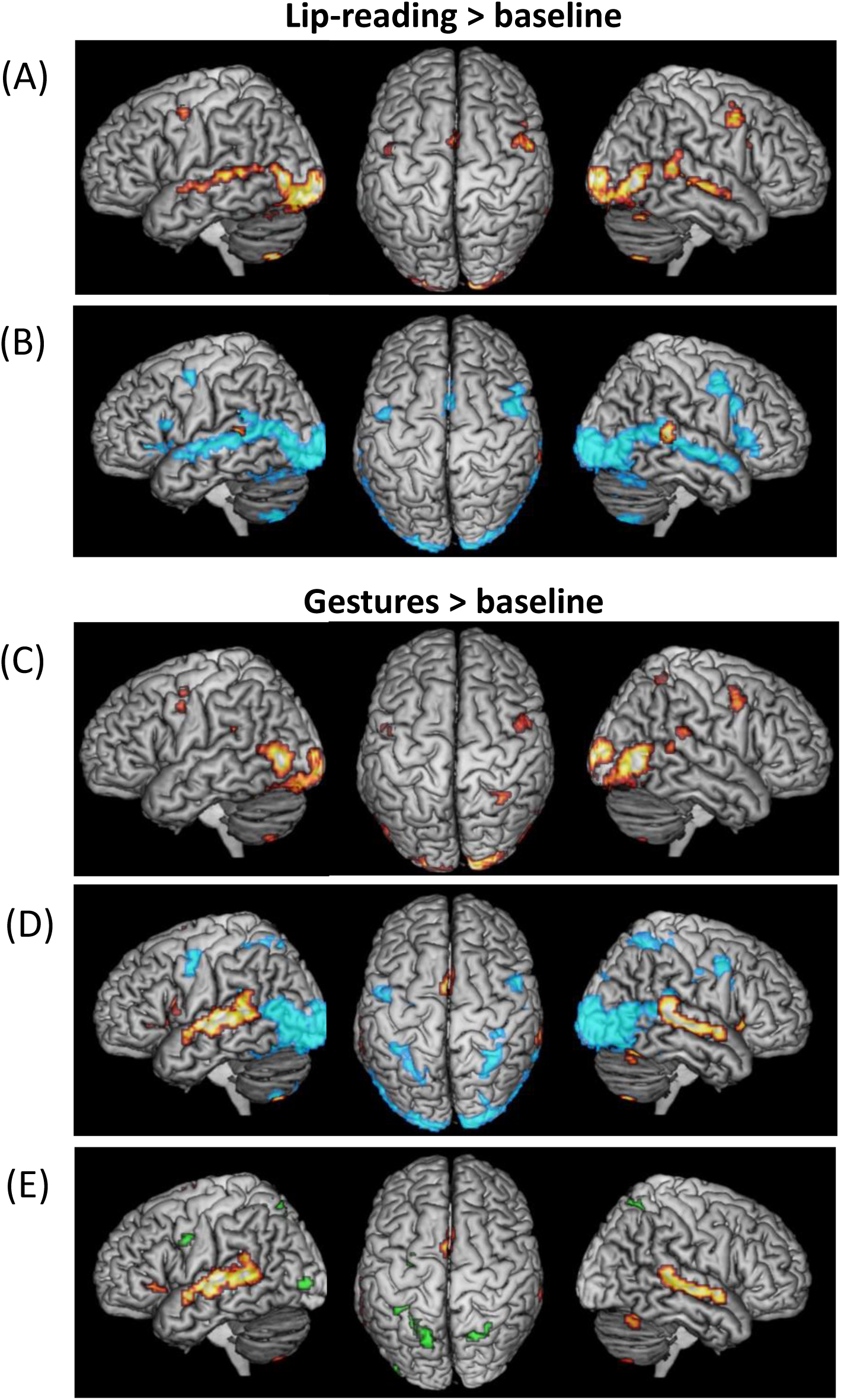
Activation by Lip-reading and Gestures > baseline. (A) Lip-reading > baseline: Activations common to the three groups; (B) Lip-reading > baseline: Activations in Controls (blue) and in Deaf > Controls (hot); (C) Gestures > baseline: Activations common to the three groups; (D) Gestures > baseline: Activations in Controls (blue) and in Deaf > Controls (hot); (E) Gestures > baseline: Activations in Deaf > Hearing (hot) and Hearing > Deaf (green).

##### Lip-reading

The conjunction of Lip-reading > baseline across the three groups activated a large subset of the regions activated by Sentences > baseline: the bilateral occipital poles, inferior occipital cortex, lateral occipital cortex, and fusiform gyrus; plus the bilateral STS/STG, precentral gyrus and SMA (Figure 3A). Note that all 3 groups also showed activation in the left IFG, with only partial overlap, explaining why it did not survive in the conjunction analysis. Pairwise comparisons across groups showed stronger activation in Deaf > Controls (masked by Deaf > baseline) in the bilateral pSTG/STS (Figure 3B).

Thus, as predicted, Lip-reading activated language areas in all groups including controls, and more so in deaf participants.

##### Gestures

The conjunction of Gestures > baseline in the three groups activated the same occipital and fusiform regions as Lip-reading and Sentences, plus the bilateral posterior tip of the STG as it joins the SMG, the bilateral postcentral and precentral gyri and the right IPS (Figure 3C). In deaf participants only, Gestures induced extensive activation of language areas (Suppl figure 1A).

Accordingly, the comparisons of Deaf > Controls (masked by Deaf > baseline) showed activations along the bilateral STG/STS, in the left IFG and in the bilateral insula and SMA (Figure 3D). The comparison of Deaf > Hearing (masked by Deaf > baseline) showed the same pattern minus the left IFG and the right insula (Figure 3E).

The comparison of Hearing > Deaf (masked by Hearing > baseline) showed enhanced activations in the same bilateral IPS, left FEF, and precentral regions as observed for Sentences, plus the left inferior and middle occipital cortex (Figure 3E). Note that controls also activated the IPS/FEF regions, at a level not differing from the Deaf group. The other pairwise comparisons between groups showed no differences. Noticeably, Gestures did not activate language areas in hearing more than in controls.

Thus, as predicted, gestures activated language areas only in the two groups of CS users. Moreover, the IPS/FEF, which were absent from Lip-reading activations, were involved in the perception of gestures, more weakly in the controls and deaf groups, and more strongly in hearing users of CS.

#### Comparison between Lip-reading and Gestures

We then contrasted Lip-reading > Gestures (masked by Lip-reading > baseline), and Gestures > Lip-reading (masked by Gestures > baseline) (Suppl tables 3-4).

##### Language areas

The previous analyses showed that language areas were activated by Lip-reading in all groups, but by Gestures in CS users only. This resulted in the activation of those areas by Lip-reading > Gestures (Figure 4A) only in the hearing and control groups: in the bilateral STG/STS in both groups, plus the left precentral gyrus in hearing users, and the bilateral IFG and SMA in controls. Accordingly, the deaf group showed weaker activation than controls in the bilateral IFG, bilateral STS and left pSTG/SMG, than hearing participants in the left precentral and postcentral gyri, and than both the control and hearing groups in the bilateral SMA.

**Figure 4.**
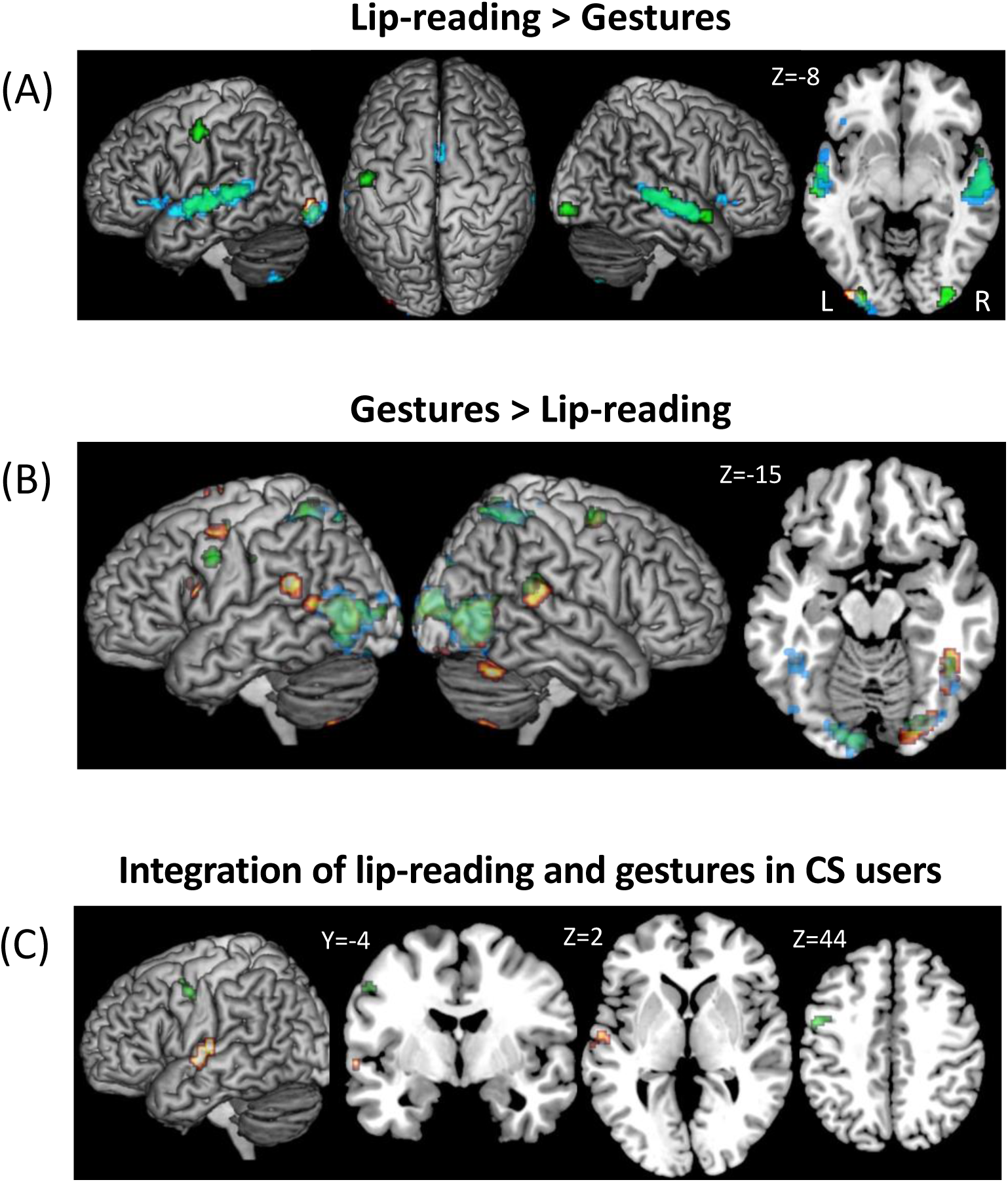
Comparison of activations by Lip-reading and Gestures. Activations in Hearing (green), in Deaf (hot) and in Control (blue) participants. (A) Activation by Lip-reading > Gestures; (B) Activation by Gestures > Lip-reading; (C) Regions integrating lip-reading and gestures in Deaf (hot) and Hearing (green) users, as defined by the conjunction of Sentences > Lip-Reading and Sentences > Gestures (both masked by Sentences > baseline in the corresponding group, thresholded at p<0.01 voxelwise).

For the opposite contrast of Gestures > Lip-reading (Figure 4B), there was no activation in language areas in controls, who had no understanding of Gestures. Conversely, deaf participants showed stronger activation to Gestures than to Lip-reading in the left IFG, right posterior middle frontal gyrus, SMA, left SMG and bilateral pSTG. Hearing participants stood in-between the other groups, with activation restricted to the left precentral gyrus and the right pSTG. The overall stronger activation by Gestures than Lip-reading in deaf participants was obvious when comparing this contrast in Deaf > Controls and in Deaf > Hearing (masked by Gestures > baseline in Deaf): both comparisons showed almost identical activation in the left IFG, the bilateral STS, the pSTG, the SMA and the insula. The Deaf > Hearing contrast additionally activated the left precentral gyrus.

##### Dorsal attentional network

The bilateral IPS-FEF, which were activated only by Gestures in the hearing users (Suppl figure 1B) and to a lesser degree in controls (Figure 3D-E), naturally appeared in the Gestures > Lip-reading contrast in those two groups (Figure 4B).

##### Visual areas

The previous analyses showed that all types of stimuli activated the same sectors of the visual system, with few differences in occipital regions. By comparing Lip-reading and Gestures, we looked for regions more engaged in the processing of one or the other type of cue.

In all 3 groups, the Lip-reading > Gestures activated a left inferior lateral occipital region, plus the symmetrical right-hemispheric region in hearing participants (Figure 4A). This activation was stronger in deaf participants than in controls.

Conversely, in all 3 groups, most sectors of the bilateral visual cortex were activated more strongly by Gestures than by Lip-reading, including the occipital poles, inferior and lateral occipital cortex and fusiform region (Figure 4B). This superiority of Gestures over Lip-reading was stronger in controls than in deaf participants (masked by Gestures > baseline in Controls) in the left lateral occipital cortex, the same region found before with the equivalent Lip-readings > Gestures contrast.

#### Integration of lip-reading and CS gestures

At the heart of the CS system is the integration of ambiguous lip-reading and gestural cues to identify unique syllables. To identify regions where this integration would take place, we used the “max criterion” of integration (Beauchamp, 2005; Ross et al., 2022). In each group, we looked for regions showing larger response for Sentences than for both Lip-reading and Gestures, restricting this analysis to regions activated during CS perception (voxelwise p<0.01). In deaf participants, this conjunction showed activation in the left mid STS (MNI -58 -11 0). In hearing users, there was activation in the left precentral gyrus (MNI -50 2 46; Figure 4C). Lowering the voxelwise threshold to p<0.01 showed additional activations in hearing users in the left IFG, and in the left SMA and anterior cingulate gyrus. Controls did not show any integrating region.

#### Activation by audible sentences

As predicted, the comparison of Audible > Silent Sentences (masked by Audible Sentences > baseline) showed no activation in deaf participants, while the conjunction of control and hearing participants (masked by the conjunction Audible > baseline of the same groups), showed massive activations in the bilateral middle temporal gyrus (MTG) and STG (Suppl figure 2A; Suppl table 5). The controls and hearing groups did not differ.

#### Activation by pseudo-sentences

Finally, we compared activations elicited by silent meaningful Sentences and by Pseudo-sentences. We first contrasted the silent meaningful Sentences > Pseudo-sentences (masked by silent meaningful Sentences > baseline). As expected, controls did not show any activation. More unexpected, deaf participants only showed a small activation cluster in the left lingual gyrus, while hearing participants showed activation in the left IFG and the bilateral left predominant insula (Suppl figure 2B; Suppl table 6). Pairwise comparisons between groups showed no significant differences.

The paucity of activations induced by this contrast is consistent with real words generating little more activation than pseudo-words (Binder et al., 2009; Taylor et al., 2013). This is because the processing of real and pseudo-language involves the same brain systems, and that pseudo-words may require additional effort than real words. However, we will later derive valuable insights into individual variability among CS users from this contrast.

#### Individual variability among hearing participants

CS mastery was variable across hearing users, while the deaf and control groups were quite homogeneous (Figure 1B). Looking for the imaging correlates of such behavioral variability, we studied the correlation of the main contrasts of interest (Sentences, Lip-reading, and Gestures > baseline, and Sentences > Pseudo-sentences, with no masking) with two individual behavioral measures: the proficiency at understanding CS sentences (as measured during the pretest), and the age of CS learning (see Suppl figure 3 for individual activation plots).

First considering the contrast of Sentences > baseline, we found a negative correlation with the participants’ age of CS learning, in the predominantly left middle cingulate and superior medial frontal gyri (Figure 5A). Furthermore, there was a negative correlation with sentence comprehension in the left middle occipital gyrus (Figure 5B). This overall effect was present separately for the comprehension of “Intermediate” and “Difficult” sentences (for the latter, activation extended to the left lingual gyrus), but not of “Easy” sentences, even when lowering the voxelwise threshold to p<0.01.

**Figure 5.**
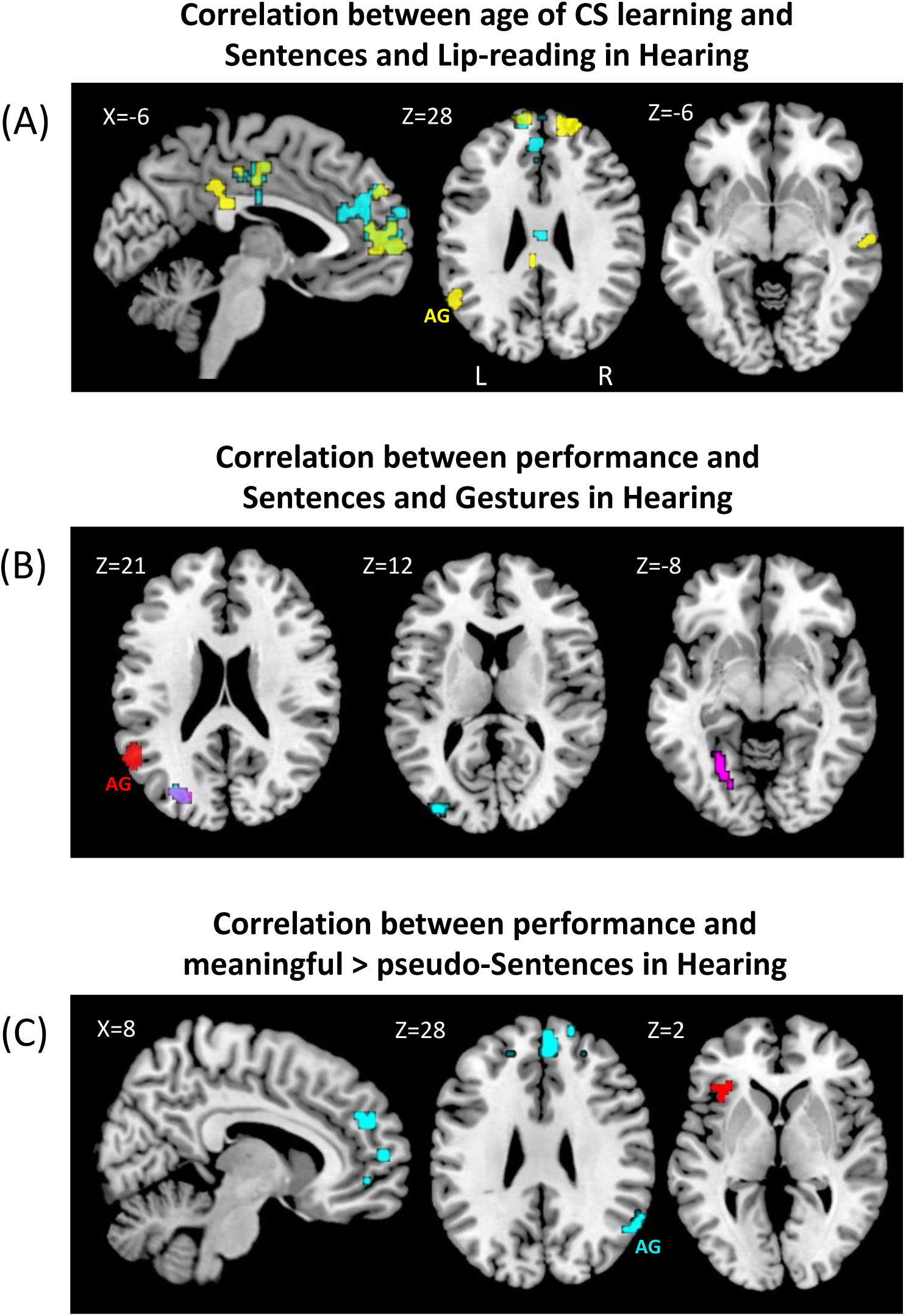
Individual variability among hearing participants. (A) Negative correlation of activations by CS Sentences (cyan) and by Lip-reading (yellow) with age of CS learning; (B) Negative correlation of activations by CS Sentences (cyan) and by Gestures (pink) with CS comprehension performance, and positive correlation of activations by Gestures (red) with CS comprehension performance; (C) Negative (cyan) and positive (red) correlation of activations by meaningful > pseudo-Sentences with CS sentences comprehension performance.

We then examined the contrast of Lip-reading > baseline. As for Sentences > baseline, there was a negative correlation with age of CS learning in the bilateral cingulate and medial superior frontal gyri, as well as in the right middle STS, the left angular gyrus and the bilateral superior and middle frontal gyri (Figure 6A). There was no correlation with sentence comprehension score.

**Figure 6.**
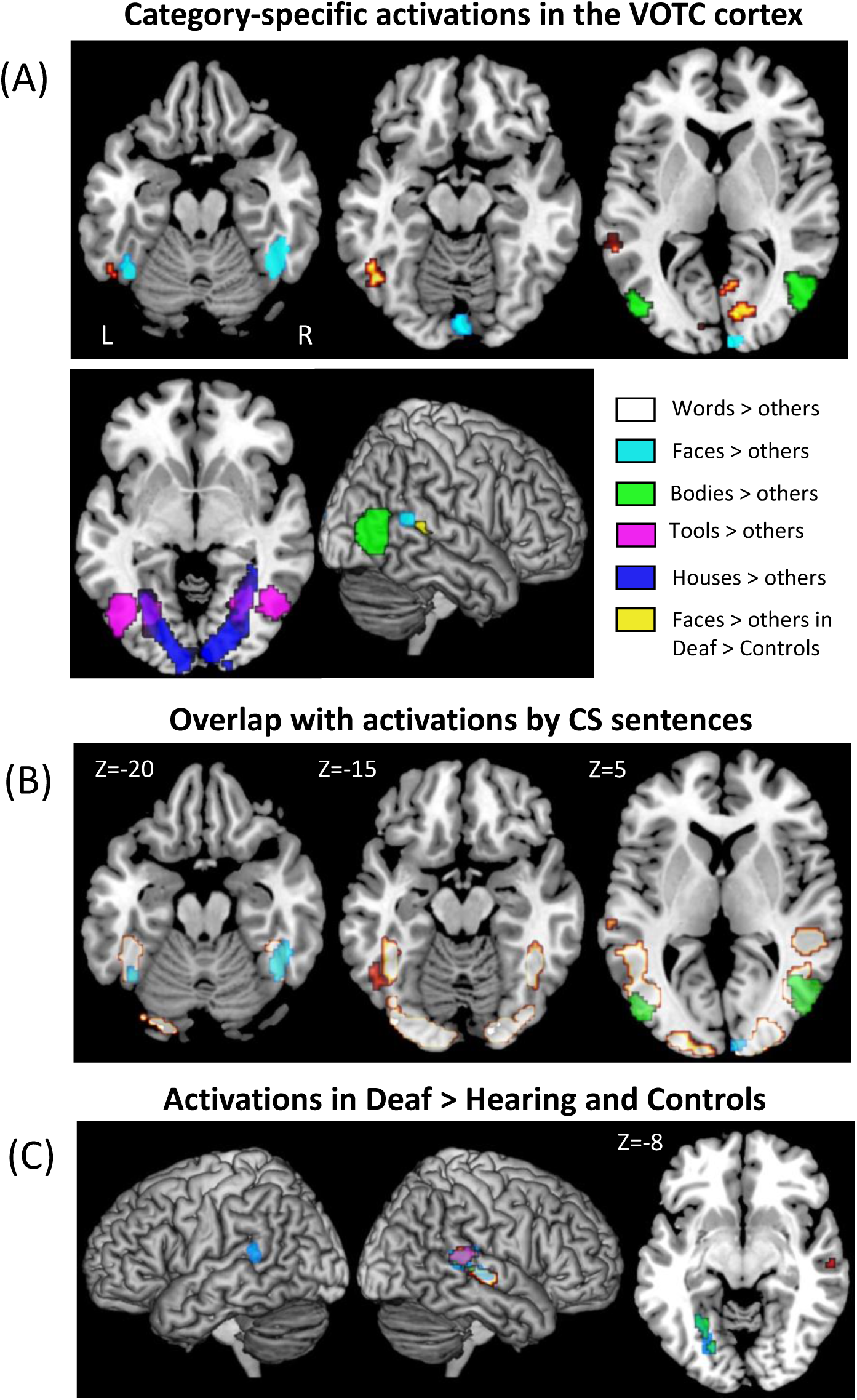
**Experiment 2**. (A) Occipitotemporal activations by each category of visual objects > all other categories, common to the three groups: Words (hot), Faces (light blue), Bodies (green), Tools (pink), Houses (dark blue). The only difference between groups was a higher selective activation by faces in Deaf than in Control participants (yellow); (B) Overlap of the category-selective activation for faces (light blue, FFA) and bodies (green, EBA), with the fusiform and lateral occipital activation by cued speech in Experiment 1 (hot outline). The word-selective VWFA (hot) was not activated by cued speech; (C) Larger activation relative to baseline, in deaf than in the average of hearing and control participants: Faces (light blue), Bodies (green), Words (hot), Tools (pink).

Turning to the Gestures > baseline contrast (Figure 5B), we found, similarly to Sentences > baseline, negative correlation with sentence comprehension in the left middle occipital gyrus and the left lingual gyrus (still absent with “Easy” sentences). Moreover, there was a positive correlation with sentence comprehension in the left angular gyrus, present with all levels of sentence difficulty (still absent with “Easy” sentences).

We then examined the contrast of meaningful Sentences > Pseudo-sentences (Figure 5C; see also Suppl figure 3C for illustration of the underlying mechanism), and found a positive correlation with sentence comprehension in the left IFG. The same result was present when examining “Difficult” sentences separately, while “Intermediate” positively correlated with the right superior medial frontal gyrus and the left cerebellum. No positive correlation was found with “Easy” sentences. Furthermore, there was a negative correlation with sentence comprehension in the right angular gyrus, the bilateral superior and superior medial frontal gyri, and the right posterior orbital frontal cortex. Examined separately, all levels of sentence difficulty yielded a negative correlation in some subset of these regions, plus the left anterior MTG and bilateral cingulate cortex for the “Difficult” sentences, and the right posterior MTG for the “Easy” sentences.

Most regions involved in those correlations belong to the so-called default mode network (DMN; Smallwood et al., 2021; Yeo et al., 2011), including the mesial prefrontal and parietal regions, the angular gyrus, and the right MTG. Overall, deactivation of the DMN diminishes as hearing users gain CS expertise, so that proficient users may require less externally oriented attention.

### Experiment 2: Brain activation during visual objects perception

In order to identify category-specific regions in the visual cortex, we presented pictures of written Words, Faces, Bodies, Tools, and Houses.

#### Behavioral results

Participants performed close to ceiling in the outlier detection task, with a mean hit rate of 97±0.04% and a mean dprime of 4.48±0.43. No significant difference in mean dprime was found between groups (all Mann-Whitney U tests p>0.05). The mean and variance of response times did not differ between groups (assessed respectively with Mann-Whitney U tests, all p>0.05, and Levene tests, all p>0.05).

#### fMRI results

##### Category-specific activations

We contrasted each category minus the average of all the others, computing the conjunction of those contrasts across the three groups. This showed the usual mosaic of occipitotemporal category-specific regions (Figure 6A; Suppl table 7). Words activated the VWFA in the left occipitotemporal sulcus, the left pSTS/STG and IFG, plus the bilateral calcarine region. Faces activated the right predominant FFA in the fusiform gyrus, the right pSTS/STG, plus the bilateral calcarine sulcus and the right cuneus. Bodies activated the Extrastriate Body Area (EBA) in the bilateral LOTC. Tools activated the bilateral collateral sulcus, the LOTC (ventral and anterior to the EBA), the right middle and superior occipital gyri, plus the bilateral IPS. Finally houses activated the PPA in the bilateral collateral sulcus and lingual gyrus, plus the right precuneus and the occipital poles.

In the conjunction of the 3 groups, the bilateral FFA (42 -51 -20 and -42 -56 -22) and EBA (50 -71 6 and -48 -78 6) overlapped respectively with the fusiform and lateral occipital activations induced by the perception of cued-speech stimuli in Experiment 1 (Figure 6B). Importantly, the VWFA, lateral to the FFA, was not activated by the perception of Sentences, Lip-reading nor Gestures. We computed the activations from Experiment 1 within spherical regions of interest (ROI) centered on individual VWFA peaks (see Methods). None of the 3 groups x 3 conditions yielded significant activation of the VWFA, except Gestures in the deaf group (t(18)=3.54, Bonferroni corrected p=0.02). There was no activation in the right-hemispheric region symmetrical to the VWFA. In contrast, spheres centered on the individual FFA were significantly activated in all groups x conditions, both in the left and right hemispheres (all Bonferroni corrected p<0.001). The same was true for the bilateral EBA (all Bonferroni corrected p<0.03, except for Lip-reading in controls in the right EBA: p=0.054).

Pairwise comparisons of category-specific activations between groups only showed increased activation in the right pSTG by Faces, in deaf participants relative to controls (masked by the same contrast in Deaf) (Figure 6A).

##### Activation relative to baseline

Beyond category-specific activation of the VOTC, pictures activated broader cortical areas relative to baseline. Pairwise comparisons showed that the control and hearing groups did not differ. The comparison of the Deaf group minus the average of the hearing and control groups (masked by activation in Deaf participants) for each visual category relative to baseline showed that deaf participants activated the right pSTS/STG more than the other participants in all categories but Houses. This activation extended more anteriorly for Words and Faces. Additional activations were present in the left collateral sulcus when viewing Faces and Bodies, and in the left pSTS/STG when viewing Faces (more posterior and smaller than in the right hemisphere) (Figure 6C; Supple table 8).

In summary, we found in all groups the usual category-selective activations in the occipitotemporal cortex. Group differences were largely restricted to the right pSTS/STG, which was more activated in deaf than in hearing and control participants for most categories of pictures. This over-activation in the Deaf group was more marked for Faces, resulting in a face-specific activation.

## Discussion

### Visual perception of cued speech

#### The role of the VWFA

Cued speech sentences activated the visual cortex almost identically across all groups, including the posterior, inferior, and lateral occipital cortex, and the fusiform region. The two latter regions overlapped with the EBA and the FFA, respectively, which is unsurprising considering that videos always featured a facing and gesturing human person (Figure 6B). More importantly, activation did not extend to the VWFA, particularly in the Deaf participants, consistent with previous findings (Aparicio et al., 2017). This is a critical finding in the debate on cortical specialization for reading in the VOTC. According to the bottom-up theory, as a result of reading acquisition, neural populations at the VWFA site become specialized for letter strings. In literate individuals, written words would be encoded by dedicated neurons, much like faces are encoded by dedicated neurons in the nearby FFA. The opposing top-down theory claims that VWFA neurons are not tuned to reading, and that reading-specific properties of the VWFA result entirely, during word reading, from top-down influences from supramodal language areas. If the only specificity of the VWFA was to grant access to language to any visual shape, one would predict activation during CS perception. This is particularly true considering that the phonological content of CS signs is comparable to the phonemic or syllabic information carried by alphabetic or syllabic scripts, which both activate the VWFA more strongly than the rest of the VOTC. The lack of VWFA activation by CS thus supports the general bottom-up view of specialization.

Moreover, Experiment 2 gave us an opportunity to compare the activation of the VWFA during word reading between deaf and hearing participants. As reviewed by Hirshorn et al. (2022), VWFA activation during reading is largely similar in deaf and in hearing persons, apart from subtle differences in the influence of phonological skills (Emmorey et al., 2016; Glezer et al., 2018). The lack of any group difference during word reading in Experiment 2 is in agreement with this overall picture.

#### The role of the LOTC

What are the visual areas that may support the processing of the gestural components of cued speech? During reading, the identification of letters must be invariant for irrelevant visual changes. There is indeed evidence that the VWFA shows invariance for case, font, and position (Cohen and Dehaene, 2004). Similarly, the identification of gestures during CS perception should be invariant for the identity of the coder, for the viewpoint, the position of the display in the visual field, etc. Parts of the bilateral LOTC show preference for the view of hands or of bodies (Bracci et al., 2010) and, using multivariate pattern analysis (MVPA), Bracci et al. (2018) found that the hand-selective regions encode hand postures in a viewpoint-invariant manner. Moreover, the LOTC cortex is also sensitive to more abstract features of gestures, such as whether they involve social interactions, and whether they involve object manipulation (Lingnau and Downing, 2015; Wurm et al., 2017; Wurm and Caramazza, 2019). Sociality is a highly relevant feature of CS gestures, due to their intrinsic communicative function. Importantly, such features of invariance and sociality overlap with all the between-groups differences which we found in the visual cortex, which were all located in the LOTC, supporting the role of those regions in implementing the visual aspects of CS expertise.

A parallel may be drawn with the functional changes occurring in the VWFA as a consequence of reading acquisition (Dehaene et al., 2015; Dehaene-Lambertz et al., 2018). Specifically, in the current study, the right-hemispheric LOTC cortex showed a reduced activation by CS Sentences in the deaf group as compared to controls (Figure 2B). Conversely in the left-hemispheric LOTC, activation by Gestures was higher in the deaf group than in controls relative to Lip-reading, a condition which never activated the LOTC in any group (Figure 4B). Aparicio et al. (2017) also found higher activation of the left LOTC during CS perception in deaf users than during a still control condition. Moreover, PPI analysis seeded on the LOTC showed that during CS perception, the left but not the right LOTC increased its connectivity with lateral temporal language regions (Aparicio et al., 2017). This overall pattern is suggestive of a shift towards the left hemisphere of the visual processing of gestures, in as much as they carry language information. This is in line with meta-analyses showing a left lateralization of such region when semantic content is carried by a perceived gesture (Cacciante et al., 2024; Yang et al., 2015). Again a parallel may be proposed with word reading, as the usual leftward lateralization of the VWFA likely results from the lateralization of language areas (Bouhali et al., 2014; Van der Haegen et al., 2012).

Moreover, the left LOTC was more activated by Gestures in hearing than in deaf users of CS (Figure 3E). First, the location of this activation further supports the role of the left LOTC in processing Gestures. Second, its high intensity may reflect the higher attentional cost of effortful deciphering in hearing CS users, congruent with the simultaneous increase in the activation of the IPS/FEF attentional network. Such a group difference was absent during the perception of full CS sentences (Figure 2C), which may reflect the higher difficulty of the task for hearing participants when viewing only Gestures, i.e. in the absence of the supporting information carried by Lip-reading. Again, a striking parallel exists with word reading: whenever readers are presented with written words degraded through tilting, letter spacing, or displacement to the visual periphery, activation increases both in the VWFA and in the bilateral IPS/FEF network (Cohen et al., 2008).

In summary, although the current design does not allow us to fully probe the functional properties of the left LOTC, this region likely implements the identification of CS gestures.

#### Fingerspelling and sign language

Although it is not the place to review the whole evidence on sign language perception, one may wonder, considering the recycling of the LOTC for CS deciphering, and of the VWFA for reading, whether those regions are also involved in the other two systems that use hand gestures as a linguistic communication channel: sign language, and its ancillary alphabetic fingerspelling. The three systems differ deeply in the function of gestures: In sign language, gestures refer to abstract linguistic entities such as morphemes, irrespective of the sound or orthographic content of their translations in spoken languages, while they denote candidate syllables in CS, and letters in fingerspelling.

The bilateral LOTC being involved in all manners of gesture perception (Yang et al., 2015), it is activated by both fingerspelling and sign language, in deaf signers and naïve controls (Emmorey et al., 2015; Liu et al., 2017; MacSweeney, 2002; Waters et al., 2007). Moreover, although the precise overlap of activations is difficult to ascertain across studies, activation by fingerspelling was stronger than by than sign language in the inferior part of the bilateral LOTC, overlapping with the LOTC region where we found sensitivity to CS knowledge, while sign language activated preferentially more dorsal LOTC regions (Emmorey et al., 2015). The VWFA also was activated by fingerspelling more than by sign language, in deaf signers only. On this basis, one may speculate (i) that ventral LOTC activation reflects the expertise in CS and fingerspelling perception, which both rely on the expert identification of static hand configurations with a language-related content; and (ii) that the VWFA is specifically attuned to the identification of letters, both printed and fingerspelled, as it is also engaged in reading novel letters shaped like faces or houses (Martin et al., 2019; Moore et al., 2014). Those hypotheses should be further assessed in appropriate within-subject studies.

#### Lip-reading

While gestures activated most of the visual cortex more strongly than lip-reading, a patch of left posterior lateral occipital cortex showed the opposite preference in all groups. This region is posterior to the face-selective FFA and OFA, and may provide input to later stages of face processing (Elbich et al., 2019). Importantly, and contrary to gestures, we found no difference between groups in the activation by Lip-reading stimuli in any part of the visual cortex. This negative finding is in keeping with the fact that this communication channel is operational in all participants. Indeed, the activation of most language areas by pure Lip-reading stimuli does not differ across groups.

#### Static images

Conducting a typical mapping of category-specific visual areas in Experiment 2, we found activation for faces in the right pSTS in deaf participants compared to the two hearing groups (Figure 6A). This over-activation has been observed in early deaf signers (Benetti et al., 2017) as well as in the context of late deafness, and it diminishes after successful cochlear implantation (Lazard and Giraud, 2017; Rouger et al., 2011). It may reflect a reassignment of the so-called “temporal voice-selective area” (Belin et al., 2000), selective for human voices in the hearing population and for face processing in deaf individuals. In both our study and Aparicio et al. (2017), this region was activated during CS perception in all three groups, with over-activation in deaf participants compared to controls and hearing users.

We also assessed non-specific activations, and found that even though the pSTS has a preference for faces, other visual categories, i.e. words, bodies and tools, elicited an over-activation in deaf participants compared to the two hearing groups (Figure 6C). Hence those findings may not be only linked to the region’s position in the secondary auditory cortex, but also to a general sensitivity to visual inputs.

Finally, static pictures of faces and bodies induced stronger activation in the left collateral sulcus of Deaf than of hearing participants (Figure 6C). The topography of this activation cluster overlaps with area V4v (Eickhoff et al., 2005; Rottschy et al., 2007). This richly connected region is thought to implement an early stage of object perception, and to guide perceptual decisions (for a review see Pasupathy et al., 2020). In deaf CS experts, for whom visual cues are the exclusive source of language perception, there is thus an increase in V4 sensitivity, specific to the two classes of objects which convey linguistic signals: human faces and bodies. Moreover, one may speculate that projection from the left V4 to the left LOTC subtending CS processing could explain the left-hemispheric lateralization of this activation.

### The Dorsal Attentional Network

Whenever the gestural component of CS was presented in Experiment 1 (Figures 2C, 3E, 4B and suppl 1B) we observed activation in the bilateral FEF and IPS. This activation was intense only in hearing CS users. It was moderate in controls, and absent in deaf participants. Frontal and parietal cortices are a source for generating attention-related signals which modulate the visual cortex from top-down (Fiebelkorn and Kastner, 2020; Mengotti et al., 2020). Among those areas, the bilateral FEF and IPS form the core of the Dorsal Attentional Network (DAN), which supports the allocation of spatial attention at locations relevant to the ongoing goals and tasks (Corbetta and Shulman, 2002). It activates for instance more during demanding than easy visual tasks, such as detecting an image in visual noise (Aben et al., 2020). Presumably, the DAN does not need to activate in the deaf participants, who are the most expert users of CS, for whom gesture identification is so automatized that it requires only minimal attentional effort. To the opposite, the hearing users of CS, who have a better experience of CS production than comprehension, require a strong attentional focus to effectively decipher the CS code. Control participants, for whom the task was too hard for additional attention to make a difference, fell in between the two other groups.

### Access to language areas

#### Commonalities across groups

With variation across groups and conditions, CS consistently activated core language areas, a set of left predominant lateral frontal and temporal regions (Fedorenko et al., 2024). Silent CS sentences elicited a set of common activations across the three groups, including controls ignorant of the CS code, in left-predominant inferior frontal, precentral, and STS/STG regions (Figure 2A). Such commonalities resulted from the shared use of lip-reading as an input code to language. Accordingly, a similar pattern of commonality was induced by pure Lip-reading stimuli (Figure 3A), but not by pure Gestures (Figure 3C). Indeed, pure Gestures induced extensive activation of language areas in deaf participants only (Figures 3D-E), which further confirms that hearing users of CS, who mastered CS less fluently than deaf participants, had a brain activation pattern in most respects comparable to the one of Controls.

#### Cued-speech expertise

Within this set of common regions, activation by CS sentences was stronger in deaf participants than in both controls (bilateral pSTS/STG and left IFG; Figure 2B), and hearing CS users (left pSTS/STG; Figure 2C). This profile nicely parallels the superior performance of deaf participants in CS comprehension as compared to naïve controls, but also, to a lesser degree, to hearing CS users (Figure 1B).

The fact that the perception of isolated gestures is sufficient to trigger linguistic activations in Deaf users may come as a surprise, as this CS component is by itself very ambiguous (∼9 possible syllables for each hand shape x position combination). Interestingly, 14 of the deaf participants declared during the debriefing having the impression of understanding at least partially these stimuli. Anecdotal testimonies exist on very proficient CS users being able to communicate using only the gestural part of CS (e.g. Weill, 2011), for example when they are talking at a distance of with their mouth full while eating. Such communications often happen in familiar contexts, so that top-down pragmatic influences certainly play an important role. There are also purely lexical constraints of the interpretation of gestures. Sequences of gestures deprived of lip-reading cues have huge numbers of potential phonological interpretations. As each gesture corresponds to ∼9 syllables, any sequence of 3 gestures may receive ∼9*9*9=729 phonological interpretations. Only a small minority of those interpretations correspond to real words, which could strongly facilitate comprehension (Ferrand et al., 2018). Still the degree of ambiguity in this linguistic input remains very high and linguistic top-down inferences seem at first insufficient for such decoding, even in a deaf population with a daily practice. Future experiments on the contribution of those factors to such performances would improve our understanding of CS processing and of the role of top-down inferences in language comprehension.

#### Language and perception

The bilateral posterior tip of the STG and precentral cortex shared an intriguing pattern of activation. They were activated by CS sentences in all groups (Figure 2A), which would be consistent with their participation in the core language system, as discussed before. Moreover, the pSTG was more activated in deaf participants than in the other groups by Sentences (Figure 2B-C), by Lip-reading (Figure 3B), and by Gestures (Figure 3D-E), suggesting that it contributed to CS expertise. However, those regions were also activated by Gestures across all groups, a surprising finding considering that Gestures carried no linguistic meaning for controls (Figure 3C-D). One possible account of this pattern is that, beyond language, both regions are also involved in action perception, in addition to the visual cortices discussed before (Wurm and Caramazza, 2022). Indeed meta-analyses show reproducible activation of the bilateral precentral gyrus and the pSTG during action observation, an activation that may even be stronger than during actual action execution (Caspers et al., 2010; Hardwick et al., 2018). Moreover strokes affecting either of those regions impair the identification of biological motion more than control moving stimuli (Saygin, 2007).

Hence, one may propose that the precentral gyrus and the pSTG act as an interface between the visual analysis of CS components and their integration in language comprehension. The same idea may apply to other visual communication systems, and indeed the left pSTG is also strongly activated in deaf signers by sign language and by fingerspelling, as compared to various controls (Emmorey et al., 2015; MacSweeney, 2002; Waters et al., 2007). More generally, this hypothesis fits well with the general role of the pSTG and the adjacent SMG whenever phonology has to be interfaced with orthography, sensorimotor processing, or lip-reading (Binder, 2017; DeWitt and Rauschecker, 2012; Hickok et al., 2018; Hickok and Poeppel, 2016; Martin et al., 2016).

### Integration of lip-reading and CS gestures

In this study, we propose that lip-reading and gestural information are integrated into CS information in the left mid STS in deaf users, and in the motor regions, mainly the left precentral gyrus, in hearing users (Figure 4C). In deaf participants, the STS location suggests that integration first occurs at the phonological level, with lip-reading and gestures information converging to form a unified phonological representation of the two components. This result runs against the alternative hypothesis that integration would occur in the visual cortex, with syllables represented as complex visual combinations of gestures and mouth configurations.

In hearing users, we found evidence of integration in motor areas, suggesting that they used a different strategy than deaf participants. Rather than relying on a unified phonological information, they relied on their good CS production skills to integrate the lips and hand movements, a process which naive controls were not able to perform. This result strengthens the idea of a precentral gyrus interfacing the visual analysis of CS components and their integration in language comprehension in expert CS producers, while the activation in controls may merely reflect the usual role of the precentral cortex in action observation. This difference in integration regions between deaf and hearing users suggests that the groups did not only differ in their proficiency in CS comprehension, but also in the very strategy which they used to process CS.

Behavioral studies indirectly suggested that CS perception may also rely on executive functions, such that each syllable would be explicitly deduced from the perceived CS components, successively taking into account gestures to constrain the interpretation of the subsequent lip movements (Attina et al., 2005). In the absence of any integration in prefrontal regions, we found no evidence to support this hypothesis (for a review see Peiffer-Smadja and Cohen, 2019).

Finally, Aparicio et al. (2017) proposed that integration occurs in the left lateral occipital cortex, based on stronger activation by full CS words than by the average of gestures and lip-reading. However, like in the present study, this region was strongly activated by full CS and by isolated gestures, and not activated by lip-reading. This pattern explains naturally the effect observed by Aparicio et al. (2017), without resorting to an integration hypothesis.

### Hearing users of cued speech

Our findings in hearing CS users may be summarized in two points. First, their behavioral mastery of CS comprehension was both substantially worse than the deaf participants’, and quite variable across individuals (Figure 1B). Second, at the group level, their pattern of brain activation was remarkably similar to the control group’s, from which it differed mostly by stronger activation of the dorsal attentional network (Figure 2B). In order to make sense of individual variability and to clarify whether hearing CS users follow a spectrum of activation patterns between controls and deaf CS experts, we studied the correlations across participants between activations on the one hand, and CS comprehension proficiency and CS age of acquisition on the other hand. Despite a relatively limited number of data points for correlation analyses, we could account for part of the individual variability.

#### Age of acquisition and the DMN

Activation by Sentences and Lip-reading was negatively correlated with the age of CS learning in left-predominant regions typical of the default mode network (DMN, Smallwood et al., 2021; Yeo et al., 2011), including mesial prefrontal and parietal regions, the angular gyrus, and the right MTG (Figure 5A). Activation in the DMN is typically suppressed whenever attention is focused on external task demands. In participants who acquired CS at an earlier age, the DMN was close to baseline, suggesting that highly automatized CS perception required little attention (Suppl figure 3A). The DMN was more deactivated in later learners, who required more focused attention to process the stimuli. This is congruent with the decrease in DMN deactivation after practicing perceptual categorization tasks (Shamloo and Helie, 2016), and with the smaller DMN deactivation in tasks that require a lower than a higher effort (Weber et al., 2022).

#### Cued speech proficiency and sentence meaning

The correlation analyses showed that the DMN was less deactivated in pseudo-compared to meaningful sentences, a result in line with previous research on the processing of spoken or written real and pseudo-words (Binder et al., 2009; Graves et al., 2017). This difference was larger in experts, as it increased with sentence comprehension scores (Figure 5C; Suppl figure 3C). Note moreover that the deactivation of the left angular gyrus (AG) component of the DMN decreased as expertise increased, a further mark that in experts CS processing required less externally oriented attention (Figure 5B).

Conversely, the left IFG was more activated in meaningful compared to pseudo sentences (Suppl figure 2B), in agreement with previous studies (Chen et al., 2023). Again, this effect of lexicality was larger in experts (Suppl figure 3C), as it increased with sentence comprehension scores, revealing a better distinction between meaningful and pseudo-sentences in the core language system of participants with better mastery of CS.

### Cued-speech proficiency and the left occipital cortex

The activation by Sentences and by Gestures was negatively correlated with CS proficiency in left ventral and lateral occipital regions (Figure 5B; Suppl figure 3B). The imaging literature on the acquisition of perceptual skills is complex, showing both increases and decreases of activation in different regions for mirror reading (e.g. Poldrack et al., 1998), or more complex inverted U-shaped curves over the period of reading acquisition (Dehaene-Lambertz et al., 2018). The current study does not allow for a decisive interpretation of this finding, which nevertheless fits with our general proposition that CS expertise rests on the functional tuning of left-hemispheric visual regions. Interestingly, the exact same left occipitotemporal patch which we identified as area V4 showed both (i) negative correlation between CS proficiency and activation by Gestures (Figure 5B), and (ii) higher activation by faces and bodies in the deaf group in Experiment 2 (Figure 6C). Despite Experiments 1 and 2 using very different stimuli and tasks, this overlap further supports the above hypothesis that the left V4 plays a role in early perceptual stages of expert CS perception.

### Open questions

We have delineated a number of mostly left-hemisphere brain regions involved in CS perception: Visual cortices where hand gestures and mouth configurations are identified, fronto-parietal regions that control visual attention in hearing CS users, distinct regions that support the integration of lip-reading and gestures depending on expertise, and core language areas where information converges. In addition, we identified correlates of individual variability among hearing CS users who differ in their CS comprehension abilities.

These findings open up a wide range of questions for future research. For example, multivariate methods would allow more direct investigation of the codes supported by each brain region, e.g., based on the decoding of gestures, mouth configurations, syllables, and their integration. Using time-resolved techniques (EEG, MEG), we could also extend this approach to the time domain, revealing the time course of CS decoding, integration, and comprehension. Beyond CS comprehension, the whole field of CS production, including the interface with language and the planning of motor commands, remains largely unexplored. Finally, CS is a valuable tool in the education of deaf children, and understanding the processes of CS learning may prove to be both fundamental and practical.

## Data availability

MRI data, behavioral data and experimental material supporting the findings of this study will be made available from the corresponding author, upon reasonable request. The stimulation, preprocessing and analysis scripts will be available on a public GitHub repository.

## Acknowledgments

We would like to thank professional CS coder Emma Andrianarison for performing all the cued speech stimuli for the videos, and Ignacio Colmenero for his central role during the recording. We are also grateful to Benoît Béranger for actively participating in the set-up of the study, and for providing us with his MRI preprocessing pipeline. We also thank Marie-Liesse Guérin and Imane Boudjelal for their assistance in the recruitment and management of participant, as well as the MRI technicians. Finally, a huge thank you to all the participants. This study would simply not have been possible without your enthusiasm and commitment.

## References

Aben, B., Buc Calderon, C., Van Den Bussche, E., Verguts, T., 2020. Cognitive Effort Modulates Connectivity between Dorsal Anterior Cingulate Cortex and Task-Relevant Cortical Areas. J. Neurosci. 40, 3838–3848. 10.1523/JNEUROSCI.2948-19.2020

Alegria, J., Charlier, B.L., Mattys, S., 1999. The Role of Lip-reading and Cued Speech in the Processing of Phonological Information in French-educated Deaf Children. European Journal of Cognitive Psychology 11, 451–472. 10.1080/095414499382255

Aparicio, M., Peigneux, P., Charlier, B., Baleriaux, D., Kavec, M., Leybaert, J., 2017. The Neural Basis of Speech Perception through Lipreading and Manual Cues: Evidence from Deaf Native Users of Cued Speech. Front Psychol 8, 426. 10.3389/fpsyg.2017.00426

Aparicio, Mario, Peigneux, P., Charlier, B., Balériaux, D., Kavec, M., Leybaert, J., 2017. The Neural Basis of Speech Perception through Lipreading and Manual Cues: Evidence from Deaf Native Users of Cued Speech. Front. Psychol. 8. 10.3389/fpsyg.2017.00426

Attina, V., Beautemps, D., Cathiard, M.-A., Odisio, M., 2004. A pilot study of temporal organization in Cued Speech production of French syllables: rules for a Cued Speech synthesizer. Speech Communication, Special Issue on Audio Visual speech processing 44, 197–214. 10.1016/j.specom.2004.10.013

Attina, V., Cathiard, M.-A., Beautemps, D., 2005. Temporal measures of hand and speech coordination during French Cued Speech production. Lecture Notes in Artificial Intelligence 3881, 13– 24.

Bayard, C., Colin, C., Leybaert, J., 2014. How is the McGurk effect modulated by Cued Speech in deaf and hearing adults? Front. Psychol. 5. 10.3389/fpsyg.2014.00416

Beauchamp, M.S., 2005. Statistical Criteria in fMRI Studies of Multisensory Integration. NI 3, 093–114. 10.1385/NI:3:2:093

Belin, P., Zatorre, R.J., Lafaille, P., Ahad, P., Pike, B., 2000. Voice-selective areas in human auditory cortex. Nature 403, 309–312. 10.1038/35002078

Benetti, S., van Ackeren, M.J., Rabini, G., Zonca, J., Foa, V., Baruffaldi, F., Rezk, M., Pavani, F., Rossion, B., Collignon, O., 2017. Functional selectivity for face processing in the temporal voice area of early deaf individuals. Proceedings of the National Academy of Sciences 114, E6437–E6446. 10.1073/pnas.1618287114

Binder, J.R., 2017. Current Controversies on Wernicke’s Area and its Role in Language. Curr Neurol Neurosci Rep 17, 58. 10.1007/s11910-017-0764-8

Binder, J.R., Desai, R.H., Graves, W.W., Conant, L.L., 2009. Where Is the Semantic System? A Critical Review and Meta-Analysis of 120 Functional Neuroimaging Studies. Cerebral Cortex 19, 2767–2796. 10.1093/cercor/bhp055

Bouhali, F., Schotten, M.T. de, Pinel, P., Poupon, C., Mangin, J.-F., Dehaene, S., Cohen, L., 2014. Anatomical Connections of the Visual Word Form Area. J. Neurosci. 34, 15402–15414. 10.1523/JNEUROSCI.4918-13.2014

Bracci, S., Caramazza, A., Peelen, M.V., 2018. View-invariant representation of hand postures in the human lateral occipitotemporal cortex. NeuroImage 181, 446–452. 10.1016/j.neuroimage.2018.07.001

Bracci, S., Ietswaart, M., Peelen, M.V., Cavina-Pratesi, C., 2010. Dissociable Neural Responses to Hands and Non-Hand Body Parts in Human Left Extrastriate Visual Cortex. Journal of Neurophysiology 103, 3389–3397. 10.1152/jn.00215.2010

Brainard, D.H., 1997. The Psychophysics Toolbox. Spatial Vision 10, 433–436. 10.1163/156856897X00357

Cacciante, L., Pregnolato, G., Salvalaggio, S., Federico, S., Kiper, P., Smania, N., Turolla, A., 2024. Language and gesture neural correlates: A meta-analysis of functional magnetic resonance imaging studies. International Journal of Language & Communication Disorders 59, 902–912. 10.1111/1460-6984.12987

Caron, C.J., Vilain, C., Schwartz, J.-L., Bayard, C., Calcus, A., Leybaert, J., Colin, C., 2023. The Effect of Cued-Speech (CS) Perception on Auditory Processing in Typically Hearing (TH) Individuals Who Are Either Naïve or Experienced CS Producers. Brain Sciences 13, 1036. 10.3390/brainsci13071036

Caspers, S., Zilles, K., Laird, A.R., Eickhoff, S.B., 2010. ALE meta-analysis of action observation and imitation in the human brain. Neuroimage 50, 1148–1167. 10.1016/j.neuroimage.2009.12.112

Chen, X., Affourtit, J., Ryskin, R., Regev, T.I., Norman-Haignere, S., Jouravlev, O., Malik-Moraleda, S., Kean, H., Varley, R., Fedorenko, E., 2023. The human language system, including its inferior frontal component in “Broca’s area,” does not support music perception. Cerebral Cortex 33, 7904–7929. 10.1093/cercor/bhad087

Cohen, L., Dehaene, S., 2004. Specialization within the ventral stream: The case for the Visual Word Form Area. Neuroimage 22, 466–476.

Cohen, L., Dehaene, S., Vinckier, F., Jobert, A., Montavont, A., 2008. Reading normal and degraded words: Contribution of the dorsal and ventral visual pathways. NeuroImage 40, 353–366. 10.1016/j.neuroimage.2007.11.036

Colin, S., Geraci, C., Leybaert, J., Petit, C., 2021. La scolarisation des élèves sourds en France : état des lieux et recommandations. Conseil scientifique de l’éducation nationale.

Corbetta, M., Shulman, G.L., 2002. Control of goal-directed and stimulus-driven attention in the brain. Nature Reviews Neuroscience 3, 201.

Cornett, R.O., 1967. CUED SPEECH. American Annals of the Deaf 112, 3–13.

Cox, R.W., 1996. AFNI: software for analysis and visualization of functional magnetic resonance neuroimages. Comput Biomed Res 29, 162–173. 10.1006/cbmr.1996.0014

Cox, R.W., Hyde, J.S., 1997. Software tools for analysis and visualization of fMRI data. NMR Biomed 10, 171–178. 10.1002/(sici)1099-1492(199706/08)10:4/5<171::aid-nbm453>3.0.co;2-l

Degano, G., Donhauser, P.W., Gwilliams, L., Merlo, P., Golestani, N., 2024. Speech prosody enhances the neural processing of syntax. Commun Biol 7, 748. 10.1038/s42003-024-06444-7

Dehaene, S., Cohen, L., Morais, J., Kolinsky, R., 2015. Illiterate to literate: behavioural and cerebral changes induced by reading acquisition. Nat Rev Neurosci 16, 234–244. 10.1038/nrn3924

Dehaene, S., Naccache, L., Cohen, L., Le Bihan, D., Mangin, J.-F., Poline, J.-B., Rivière, D., 2001. Cerebral mechanisms of word masking and unconscious repetition priming. Nat Neurosci 4, 752.

Dehaene-Lambertz, G., Monzalvo, K., Dehaene, S., 2018. The emergence of the visual word form: Longitudinal evolution of category-specific ventral visual areas during reading acquisition. PLoS Biol 16, e2004103. 10.1371/journal.pbio.2004103

Desai, S., Stickney, G., Zeng, F.-G., 2008. Auditory-visual speech perception in normal-hearing and cochlear-implant listeners. J Acoust Soc Am 123, 428–440. 10.1121/1.2816573

DeWitt, I., Rauschecker, J.P., 2012. Phoneme and word recognition in the auditory ventral stream. Proc Natl Acad Sci U S A 109, E505–14. 10.1073/pnas.1113427109

Eickhoff, S.B., Stephan, K.E., Mohlberg, H., Grefkes, C., Fink, G.R., Amunts, K., Zilles, K., 2005. A new SPM toolbox for combining probabilistic cytoarchitectonic maps and functional imaging data. Neuroimage 25, 1325–35. 10.1016/j.neuroimage.2004.12.034

Elbich, D.B., Molenaar, P.C.M., Scherf, K.S., 2019. Evaluating the organizational structure and specificity of network topology within the face processing system. Human Brain Mapping 40, 2581– 2595. 10.1002/hbm.24546

Emmorey, K., McCullough, S., Weisberg, J., 2016. The neural underpinnings of reading skill in deaf adults. Brain and language 160, 11–20.

Emmorey, K., McCullough, S., Weisberg, J., 2015. Neural correlates of fingerspelling, text, and sign processing in deaf American Sign Language–English bilinguals. Language, Cognition and Neuroscience 30, 749–767. 10.1080/23273798.2015.1014924

Fedorenko, E., Ivanova, A.A., Regev, T.I., 2024. The language network as a natural kind within the broader landscape of the human brain. Nat. Rev. Neurosci. 25, 289–312. 10.1038/s41583-024-00802-4

Fenlon, J., Cormier, K., Brentari, D., 2017. The phonology of sign languages, in: The Routledge Handbook of Phonological Theory. Routledge.

Ferrand, L., Méot, A., Spinelli, E., New, B., Pallier, C., Bonin, P., Dufau, S., Mathôt, S., Grainger, J., 2018. MEGALEX: A megastudy of visual and auditory word recognition. Behav Res 50, 1285–1307. 10.3758/s13428-017-0943-1

Fiebelkorn, I.C., Kastner, S., 2020. Functional Specialization in the Attention Network. Annu. Rev. Psychol. 71, 221–249. 10.1146/annurev-psych-010418-103429

Friedmann, N., Rusou, D., 2015. Critical period for first language: the crucial role of language input during the first year of life. Current Opinion in Neurobiology, Circuit plasticity and memory 35, 27–34. 10.1016/j.conb.2015.06.003

Gardiner-Walsh, S.J., Giese, K., Walsh, T.P., 2020. Cued Speech: Evolving Evidence 1968–2018. Deafness & Education International 0, 1–22. 10.1080/14643154.2020.1755144

Gaser, C., 2020. CAT12.

Glezer, L.S., Weisberg, J., Farnady, C.O., McCullough, S., Midgley, K.J., J Holcomb, P., Emmorey, K., 2018. Orthographic and phonological selectivity across the reading system in deaf skilled readers. Neuropsychologia 117, 500–512. 10.1016/j.neuropsychologia.2018.07.010

Graves, W.W., Boukrina, O., Mattheiss, S.R., Alexander, E.J., Baillet, S., 2017. Reversing the Standard Neural Signature of the Word–Nonword Distinction. Journal of Cognitive Neuroscience 29, 79–94. 10.1162/jocn_a_01022

Hardwick, R.M., Caspers, S., Eickhoff, S.B., Swinnen, S.P., 2018. Neural correlates of action: Comparing meta-analyses of imagery, observation, and execution. Neuroscience & Biobehavioral Reviews 94, 31–44. 10.1016/j.neubiorev.2018.08.003

Hickok, G., Poeppel, D., 2016. Chapter 25 - Neural Basis of Speech Perception, in: Hickok, G., Small, S.L. (Eds.), Neurobiology of Language. Academic Press, San Diego, pp. 299–310. 10.1016/B978-0-12-407794-2.00025-0

Hickok, G., Rogalsky, C., Matchin, W., Basilakos, A., Cai, J., Pillay, S., Ferrill, M., Mickelsen, S., Anderson, S.W., Love, T., Binder, J., Fridriksson, J., 2018. Neural Networks Supporting Audiovisual Integration for Speech: A Large-Scale Lesion Study. Cortex 103, 360–371. 10.1016/j.cortex.2018.03.030

Hirshorn, E.A., Dye, M.W., Hauser, P.C., Supalla, T., Bavelier, D., 2022. Reading in Deaf Individuals: Examining the role of visual word form area. Changing Brains 117–137.

International Academy Supporting Adaptations of Cued Speech (AISAC) [WWW Document], 2020. . AISAC. URL https://www.academieinternationale.org/list-of-cued-languages (accessed 11.25.24).

Kasper, L., Bollmann, S., Diaconescu, A.O., Hutton, C., Heinzle, J., Iglesias, S., Hauser, T.U., Sebold, M., Manjaly, Z.-M., Pruessmann, K.P., Stephan, K.E., 2017. The PhysIO Toolbox for Modeling Physiological Noise in fMRI Data. Journal of Neuroscience Methods 276, 56–72. 10.1016/j.jneumeth.2016.10.019

Lazard, D.S., Giraud, A.-L., 2017. Faster phonological processing and right occipito-temporal coupling in deaf adults signal poor cochlear implant outcome. Nat Commun 8, 14872. 10.1038/ncomms14872

Lingnau, A., Downing, P.E., 2015. The lateral occipitotemporal cortex in action. Trends in Cognitive Sciences 19, 268–277. 10.1016/j.tics.2015.03.006

Liu, L., Yan, X., Liu, J., Xia, M., Lu, C., Emmorey, K., Chu, M., Ding, G., 2017. Graph theoretical analysis of functional network for comprehension of sign language. Brain Res 1671, 55–66. 10.1016/j.brainres.2017.06.031

MacSweeney, M., 2002. Neural systems underlying British Sign Language and audio-visual English processing in native users. Brain 125, 1583–1593. 10.1093/brain/awf153

MacSweeney, M., Capek, C.M., Campbell, R., Woll, B., 2008. The signing brain: the neurobiology of sign language. Trends in Cognitive Sciences 12, 432–440. 10.1016/j.tics.2008.07.010

Martin, A., Kronbichler, M., Richlan, F., 2016. Dyslexic brain activation abnormalities in deep and shallow orthographies: A meta-analysis of 28 functional neuroimaging studies 24.

Martin, L., Durisko, C., Moore, M.W., Coutanche, M.N., Chen, D., Fiez, J.A., 2019. The VWFA Is the Home of Orthographic Learning When Houses Are Used as Letters. eNeuro 6. 10.1523/ENEURO.0425-17.2019

Mengotti, P., Käsbauer, A.-S., Fink, G.R., Vossel, S., 2020. Lateralization, functional specialization, and dysfunction of attentional networks. Cortex 132, 206–222. 10.1016/j.cortex.2020.08.022

Moore, M.W., Durisko, C., Perfetti, C.A., Fiez, J.A., 2014. Learning to Read an Alphabet of Human Faces Produces Left-lateralized Training Effects in the Fusiform Gyrus. Journal of Cognitive Neuroscience 26, 896–913. 10.1162/jocn_a_00506

Nakamura, K., Dehaene, S., Jobert, A., Le Bihan, D., Kouider, S., 2005. Subliminal convergence of Kanji and Kana words: further evidence for functional parcellation of the posterior temporal cortex in visual word perception. J Cogn Neurosci 17, 954–68.

Oldfield, R.C., 1971. The assessment and analysis of handedness: The Edinburgh inventory. Neuropsychologia 9, 97–113. 10.1016/0028-3932(71)90067-4

Pasupathy, A., Popovkina, D.V., Kim, T., 2020. Visual Functions of Primate Area V4. Annu Rev Vis Sci 6, 363–385. 10.1146/annurev-vision-030320-041306

Peiffer-Smadja, N., Cohen, L., 2019. The cerebral bases of the bouba-kiki effect. NeuroImage 186, 679–689. 10.1016/j.neuroimage.2018.11.033

Poldrack, R.A., Desmond, J.E., Glover, G.H., Gabrieli, J.D., 1998. The neural basis of visual skill learning: an fMRI study of mirror reading. Cereb Cortex 8, 1–10.

Price, C.J., Devlin, J.T., 2011. The interactive account of ventral occipitotemporal contributions to reading. Trends Cogn Sci 15, 246–53. 10.1016/j.tics.2011.04.001

Ross, L.A., Molholm, S., Butler, J.S., Bene, V.A.D., Foxe, J.J., 2022. Neural correlates of multisensory enhancement in audiovisual narrative speech perception: A fMRI investigation. NeuroImage 263, 119598. 10.1016/j.neuroimage.2022.119598

Rottschy, C., Eickhoff, S.B., Schleicher, A., Mohlberg, H., Kujovic, M., Zilles, K., Amunts, K., 2007. Ventral visual cortex in humans: cytoarchitectonic mapping of two extrastriate areas. Hum Brain Mapp 28, 1045–59. 10.1002/hbm.20348

Rouger, J., Lagleyre, S., Démonet, J., Fraysse, B., Deguine, O., Barone, P., 2011. Evolution of crossmodal reorganization of the voice area in cochlear-implanted deaf patients. Hum Brain Mapp 33, 1929–1940. 10.1002/hbm.21331

Rueckl, J.G., Paz-Alonso, P.M., Molfese, P.J., Kuo, W.-J., Bick, A., Frost, S.J., Hancock, R., Wu, D.H., Mencl, W.E., Duñabeitia, J.A., Lee, J.-R., Oliver, M., Zevin, J.D., Hoeft, F., Carreiras, M., Tzeng, O.J.L., Pugh, K.R., Frost, R., 2015. Universal brain signature of proficient reading: Evidence from four contrasting languages. Proceedings of the National Academy of Sciences 112, 15510–15515. 10.1073/pnas.1509321112

Saygin, A.P., 2007. Superior temporal and premotor brain areas necessary for biological motion perception. Brain 130, 2452–2461. 10.1093/brain/awm162

Schorr, E.A., Fox, N.A., Wassenhove, V. van, Knudsen, E.I., 2005. Auditory-visual fusion in speech perception in children with cochlear implants. PNAS 102, 18748–18750. 10.1073/pnas.0508862102

Shamloo, F., Helie, S., 2016. Changes in default mode network as automaticity develops in a categorization task. Behavioural Brain Research 313, 324–333. 10.1016/j.bbr.2016.07.029

Smallwood, J., Bernhardt, B.C., Leech, R., Bzdok, D., Jefferies, E., Margulies, D.S., 2021. The default mode network in cognition: a topographical perspective. Nat Rev Neurosci 22, 503–513. 10.1038/s41583-021-00474-4

Taylor, J.S.H., Rastle, K., Davis, M.H., 2013. Can cognitive models explain brain activation during word and pseudoword reading? A meta-analysis of 36 neuroimaging studies. Psychological Bulletin 139, 766–791. 10.1037/a0030266

The tedana Community, Ahmed, Z., Bandettini, P.A., Bottenhorn, K.L., Caballero-Gaudes, C., Dowdle, L.T., DuPre, E., Gonzalez-Castillo, J., Handwerker, D., Heunis, S., Kundu, P., Laird, A.R., Markello, R., Markiewicz, C.J., Maullin-Sapey, T., Moia, S., Salo, T., Staden, I., Teves, J., Uruñuela, E., Vaziri-Pashkam, M., Whitaker, K., 2021. ME-ICA/tedana: 0.0.10. 10.5281/zenodo.4725985

Trettenbrein, P.C., Papitto, G., Friederici, A.D., Zaccarella, E., 2021. Functional neuroanatomy of language without speech: An ALE meta-analysis of sign language. Human Brain Mapping 42, 699– 712. 10.1002/hbm.25254

Trezek, B.J., 2017. Cued Speech and the Development of Reading in English: Examining the Evidence. J Deaf Stud Deaf Educ 22, 349–364. 10.1093/deafed/enx026

Van der Haegen, L., Cai, Q., Brysbaert, M., 2012. Colateralization of Broca’s area and the visual word form area in left-handers: fMRI evidence. Brain Lang 122, 171–8. 10.1016/j.bandl.2011.11.004 S0093-934X(11)00186-6 [pii]

Waters, D., Campbell, R., Capek, C.M., Woll, B., David, A.S., McGuire, P.K., Brammer, M.J., MacSweeney, M., 2007. Fingerspelling, signed language, text and picture processing in deaf native signers: The role of the mid-fusiform gyrus. NeuroImage 35, 1287–1302. 10.1016/j.neuroimage.2007.01.025

Weber, S., Aleman, A., Hugdahl, K., 2022. Involvement of the default mode network under varying levels of cognitive effort. Sci Rep 12, 6303. 10.1038/s41598-022-10289-7

Weill, A.-L., 2011. Les trois heureux paradoxes de la Langue française Parlée Completée [The three lucky paradoxes of French Cued Speech], in: La Langue Française Parlée Completée (LPC): Fondements et Perspectives [French Cued Speech: Principles and Perspectives]. Solal, Marseille, pp. 87–95.

Werker, J.F., Hensch, T.K., 2015. Critical Periods in Speech Perception: New Directions. Annual Review of Psychology 66, 173–196. 10.1146/annurev-psych-010814-015104

Wurm, M.F., Caramazza, A., 2022. Two ‘what’ pathways for action and object recognition. Trends in Cognitive Sciences 26, 103–116. 10.1016/j.tics.2021.10.003

Wurm, M.F., Caramazza, A., 2019. Lateral occipitotemporal cortex encodes perceptual components of social actions rather than abstract representations of sociality. NeuroImage 202, 116153. 10.1016/j.neuroimage.2019.116153

Wurm, M.F., Caramazza, A., Lingnau, A., 2017. Action Categories in Lateral Occipitotemporal Cortex Are Organized Along Sociality and Transitivity. J. Neurosci. 37, 562–575. 10.1523/JNEUROSCI.1717-16.2016

Yang, J., Andric, M., Mathew, M.M., 2015. The neural basis of hand gesture comprehension: A meta-analysis of functional magnetic resonance imaging studies. Neuroscience & Biobehavioral Reviews 57, 88–104. 10.1016/j.neubiorev.2015.08.006

Yeo, B.T., Krienen, F.M., Sepulcre, J., Sabuncu, M.R., Lashkari, D., Hollinshead, M., Roffman, J.L., Smoller, J.W., Zöllei, L., Polimeni, J.R., 2011. The organization of the human cerebral cortex estimated by intrinsic functional connectivity. Journal of neurophysiology.

Zhan, M., Pallier, C., Agrawal, A., Dehaene, S., Cohen, L., 2023. Does the visual word form area split in bilingual readers? A millimeter-scale 7-T fMRI study. Science Advances 9, eadf6140.

