## Supplementary figures for "Seeing speech: neural mechanisms of cued speech perception in prelingually deaf and hearing users"

**Activation by Gestures > baseline in Deaf**

(A)

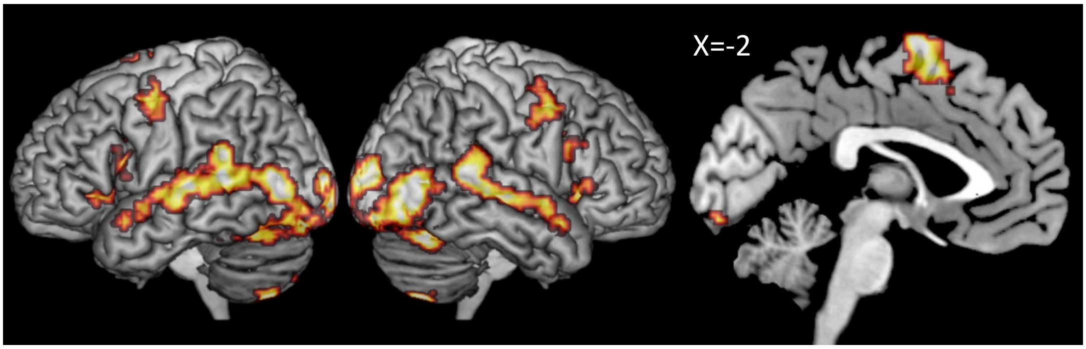


**Activation by Gestures > baseline in Hearing**

(B)

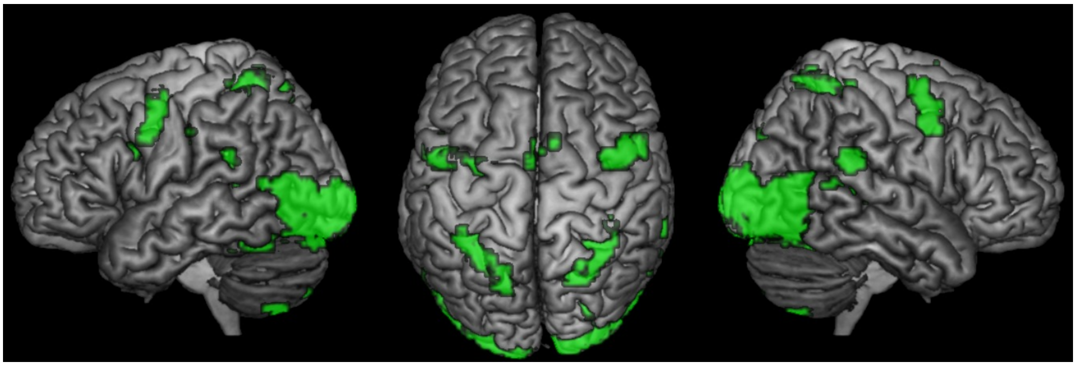

**Supplementary figure 1.** (A) Activation by Gestures > baseline in deaf users; (B) Activation by Gestures > baseline in hearing users.

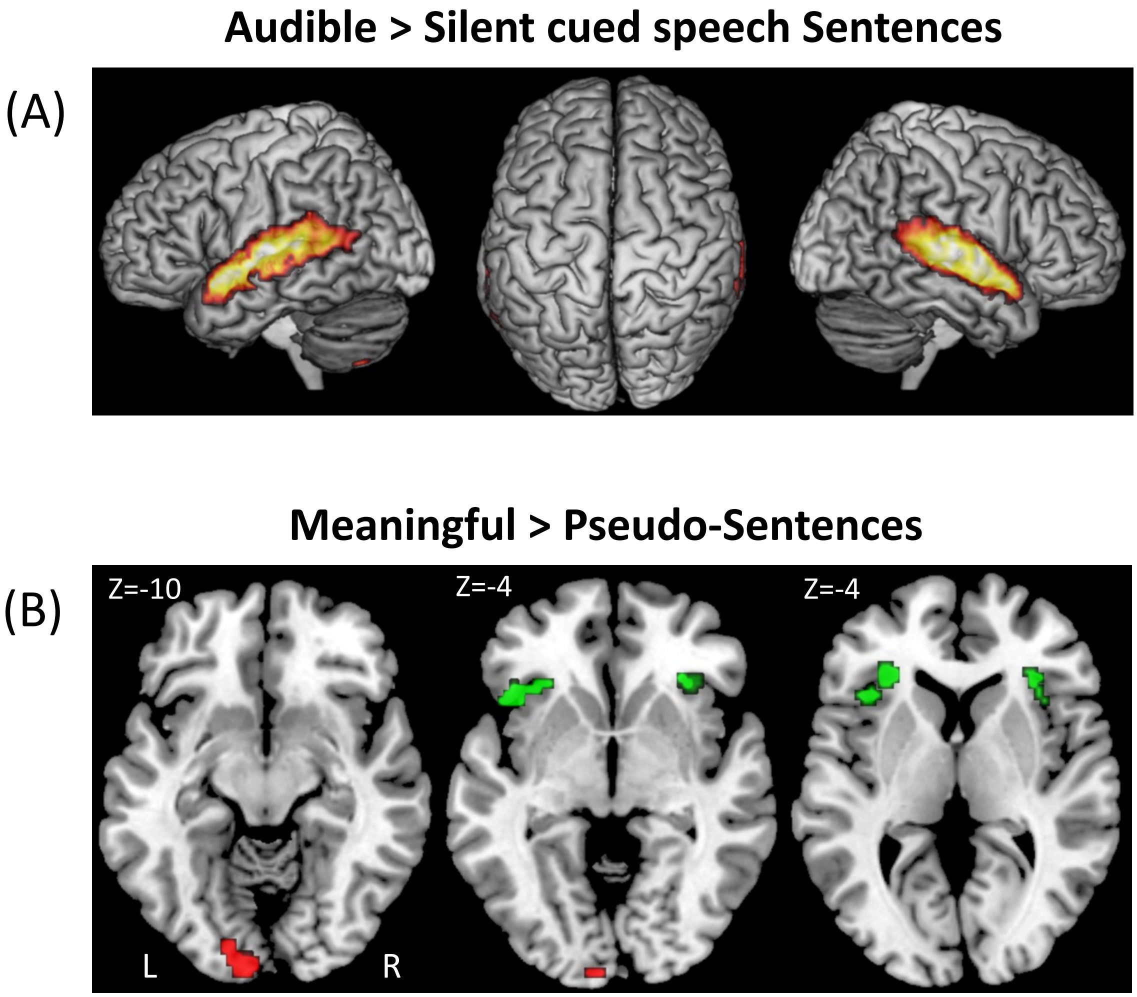


**Supplementary figure 2.** (A) Activation by Audible > silent Sentences : Activations common to hearing and control participants; (B) Activation by meaningful > Pseudo-sentences in Deaf (red) and Hearing (green) participants.

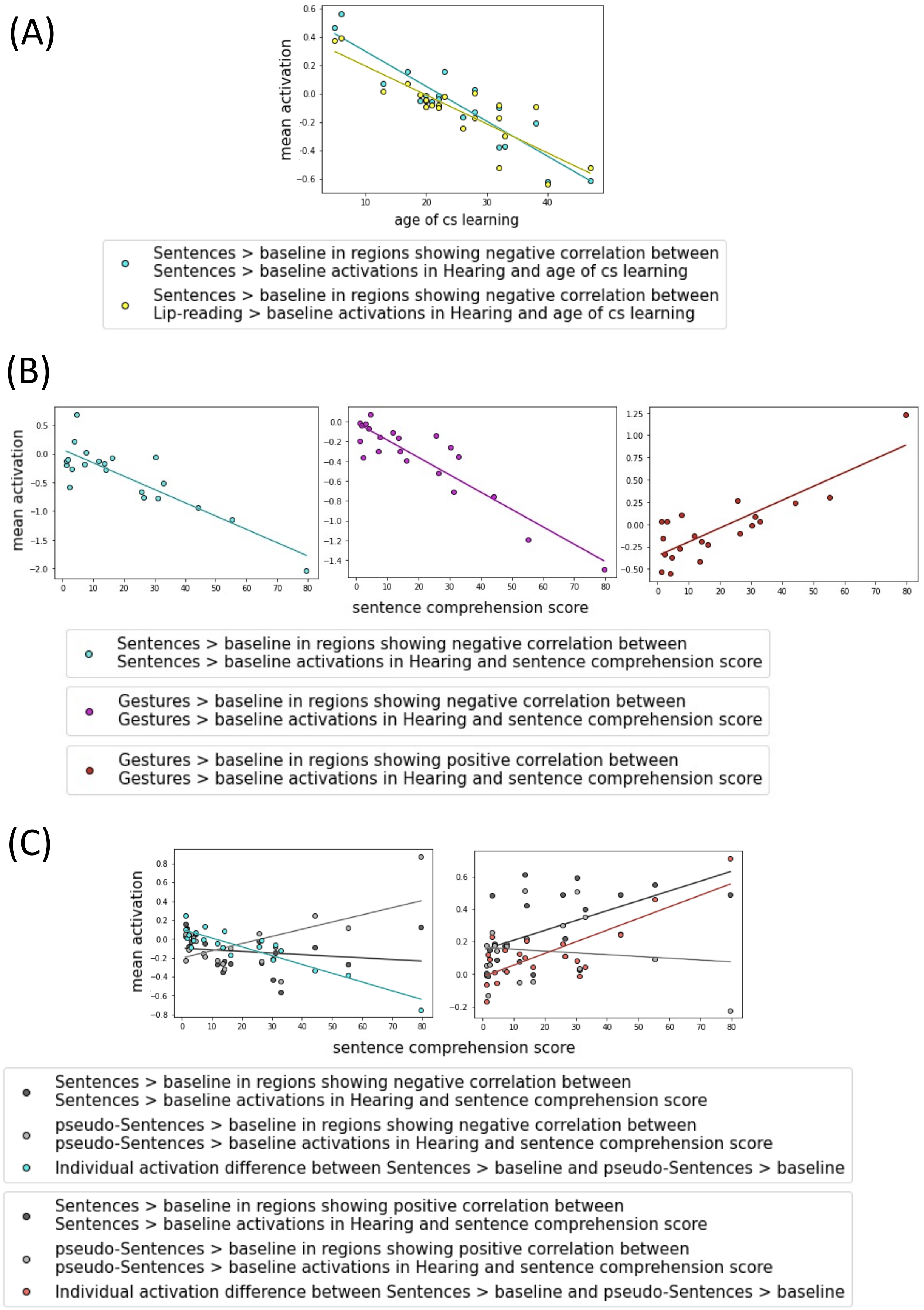


**Supplementary figure 3 – Plots of individual activation in hearing users, showing the links with the age of CS acquisition and the CS comprehension proficiency.** Order of display and color codes are identical to corresponding figure 5 activations.
