## Supplementary tables for "Seeing speech: neural mechanisms of cued speech perception in prelingually deaf and hearing users"

| Region | Conjunction 3 groups | | | | Deaf > Controls | | | |
|---|---|---|---|---|---|---|---|---|
| | x | y | z | Z | x | y | z | Z |
| L Calcarine sulcus | -15 | -101 | -7 | 7.4 | | | | |
| R Calcarine sulcus | 15 | -101 | -2 | 7.18 | | | | |
| L Inferior occipital gyrus | -12 | -91 | -14 | 6.33 | | | | |
| R Inferior occipital gyrus | 12 | -91 | -10 | 6.21 | | | | |
| L Middle occipital gyrus | -20 | -96 | -2 | 7.1 | | | | |
| L LOTC | -45 | -71 | 6 | 5.75 | | | | |
| R LOTC | 50 | -68 | -20 | 6.73 | | | | |
| L Fusiform gyrus | -42 | -51 | -22 | 5.39 | | | | |
| R Fusiform gyrus | 42 | -48 | -20 | 5.13 | | | | |
| L aSTS/STG | -60 | -18 | -4 | 4.04 | | | | |
| R aSTS/STG | 50 | -38 | 8 | 6.51 | | | | |
| L mSTS/STG | -65 | -28 | 3 | 4.24 | | | | |
| R mSTS/STG | 55 | -26 | 0 | 5.4 | | | | |
| L pSTS/STG | -50 | -46 | 8 | 5.11 | -58 | -41 | 10 | 4.51 |
| | | | | | -45 | -38 | 18 | 3.65 |
| R pSTS/STG | 55 | -16 | -7 | 5.37 | 62 | -36 | 8 | 4.79 |
| L Precentral gyrus | -42 | -1 | 50 | 3.56 | | | | |
| R Precentral gyrus | 48 | 2 | 46 | 4.39 | | | | |
| L/R SMA | -2 | 6 | 58 | 4.55 | | | | |
| L Cerebellum | -28 | -66 | -54 | 5.12 | | | | |
| | -10 | -74 | -44 | 4.3 | | | | |
| | -30 | -61 | -27 | 4.53 | | | | |
| | -15 | -71 | -24 | 4.14 | | | | |
| | -42 | -64 | -27 | 3.75 | | | | |
| R Cerebellum | 42 | -64 | -27 | 4.68 | | | | |
| | 32 | -61 | -24 | 4.34 | | | | |
| | 30 | -64 | -52 | 5.29 | | | | |

**Supplementary table 1 - Lip-reading > baseline.** Hemisphere and anatomical regions, MNI coordinates and Z score of peak activations. Voxelwise threshold p<0.001, clusterwise threshold p<0.05 FDR-corrected over the whole brain. Comparisons between groups are masked by the considered activation in the first group. L=left; R=right; a=anterior; m=middle; p=posterior; LOTC=lateral occipitotemporal cortex; STS=superior temporal sulcus; STG=superior temporal gyrus; SMA=supplementary motor area

| Region | Conjunction 3 groups | | | | Deaf | | | | Controls | | | | Deaf > Controls | | | | Deaf > Hearing | | | | Hearing > Deaf | | | |
|---|---|---|---|---|---|---|---|---|---|---|---|---|---|---|---|---|---|---|---|---|---|---|---|---|
| | x | y | z | Z | x | y | z | Z | x | y | z | Z | x | y | z | Z | x | y | z | Z | x | y | z | Z |
| L Calcarine sulcus | -12 | -101 | -7 | 7.37 | -15 | -98 | -2 | 6.2 | -8 | -101 | -7 | 6.5 | | | | | | | | | | | | |
| R Calcarine sulcus | 12 | -101 | 3 | > 8 | 10 | -101 | 3 | 6.78 | 12 | -101 | 3 | 6.44 | | | | | | | | | | | | |
| L Inferior occipital gyrus | -22 | -98 | 3 | 6.52 | | | | | | | | | | | | | | | | | -40 | -71 | -10 | 3.63 |
| L Middle occipital gyrus | | | | | | | | | | | | | | | | | | | | | -38 | -86 | 0 | 3.95 |
| R Superior occipital gyrus | | | | | | | | | 28 | -71 | 30 | 4.19 | | | | | | | | | | | | |
| L LOTC | -48 | -71 | 3 | 7.22 | -48 | -71 | 3 | 6.05 | -45 | -71 | 3 | 6.47 | | | | | | | | | | | | |
| R LOTC | 48 | -68 | -2 | > 8 | 50 | -68 | -4 | 6.4 | 42 | -64 | 3 | 6.25 | | | | | | | | | | | | |
| R Fusiform gyrus | 42 | -56 | -17 | 5.49 | | | | | 42 | -48 | -22 | 6.3 | | | | | | | | | | | | |
| L aSTS/STG | | | | | -52 | 12 | -17 | 4.06 | | | | | -62 | -18 | -2 | 5.62 | | | | | | | | |
| R aSTS/STG | | | | | | | | | | | | | 60 | -16 | -4 | 5.56 | 60 | 2 | -4 | 6.04 | | | | |
| L mSTS/STG | | | | | | | | | | | | | -65 | -24 | 6 | 5.63 | -65 | -21 | 3 | 6.38 | | | | |
| R mSTS/STG | | | | | 58 | -16 | -4 | 5.49 | | | | | 60 | -28 | 0 | 5.97 | 60 | -16 | -4 | 5.69 | | | | |
| L pSTS/STG | | | | | -60 | -41 | 10 | 6.03 | | | | | -58 | -41 | 10 | 5.99 | -58 | -34 | 6 | 5.7 | | | | |
| R pSTS/STG | 50 | -38 | 8 | 4.21 | 52 | -36 | 3 | 5.66 | | | | | 62 | -36 | 8 | 6.11 | 65 | -36 | 8 | 5.51 | | | | |
| L MTS/MTG | -48 | -44 | 10 | 4.03 | | | | | | | | | | | | | | | | | | | | |
| L Supramarginal gyrus | -45 | -41 | 26 | 4.31 | | | | | -45 | -38 | 26 | 4.29 | | | | | | | | | | | | |
| R Supramarginal gyrus | 60 | -36 | 23 | 4.28 | | | | | | | | | | | | | | | | | | | | |
| L IFG (pars opercularis) | | | | | -48 | 12 | 16 | 4.82 | 48 | 12 | 23 | 3.71 | -48 | 12 | 13 | 4.59 | | | | | | | | |
| L IFG (pars orbitalis) | | | | | -40 | 24 | 0 | 4.41 | | | | | | | | | | | | | | | | |
| R IFG (pars opercularis) | | | | | 52 | 16 | 23 | 3.97 | | | | | | | | | | | | | | | | |
| R IFG (pars triangularis) | | | | | 52 | 19 | 0 | 3.81 | | | | | | | | | | | | | | | | |
| L Insula | | | | | -30 | 16 | 3 | 4.58 | | | | | -35 | 26 | -2 | 3.98 | -45 | 26 | -4 | 3.99 | | | | |
| R Insula | | | | | 35 | 22 | -2 | 5.01 | | | | | 35 | 26 | 6 | 4.58 | | | | | | | | |
| L Precentral gyrus | -42 | -4 | 50 | 4.03 | -42 | -4 | 53 | 5.28 | -45 | -4 | 50 | 4.49 | | | | | | | | | -52 | 4 | 33 | 4.18 |
| | | | | | | | | | -48 | -4 | 23 | 3.53 | | | | | | | | | | | | |
| R Precentral gyrus | 48 | 4 | 43 | 4.15 | 45 | 6 | 43 | 5.22 | 48 | 6 | 46 | 4.36 | | | | | | | | | | | | |
| R Postcentral gyrus | 35 | -34 | 53 | 4.26 | 32 | -36 | 50 | 3.86 | 35 | -34 | 53 | 4.93 | | | | | | | | | | | | |
| | | | | | | | | | 50 | -16 | 36 | 4.26 | | | | | | | | | | | | |
| L/R SMA | | | | | -2 | 4 | 68 | 6.29 | | | | | -2 | 2 | 68 | 4.99 | -2 | 4 | 68 | 5.09 | | | | |
| L IPS | | | | | | | | | -18 | -56 | 53 | 4.73 | | | | | | | | | -38 | -36 | 40 | 5.35 |
| | | | | | | | | | -45 | -24 | 38 | 4.37 | | | | | | | | | -20 | -61 | 60 | 4.83 |
| R IPS | 32 | -51 | 60 | 4.25 | | | | | 32 | -54 | 63 | 4.81 | | | | | | | | | 15 | -64 | 60 | 3.66 |
| L FEF | | | | | | | | | -28 | -6 | 48 | 4.3 | | | | | | | | | -25 | -6 | 60 | 4.55 |
| R FEF | | | | | | | | | | | | | | | | | | | | | | | | |
| L Cerebellum | -10 | -74 | -44 | 5 | -25 | -61 | -44 | 5.26 | -10 | -76 | -44 | 5.32 | | | | | | | | | | | | |
| | | | | | -10 | -81 | -44 | 4.24 | -28 | -64 | -50 | 4.36 | | | | | | | | | | | | |
| | | | | | -15 | -74 | -22 | 4.85 | | | | | | | | | | | | | | | | |
| R Cerebellum | 12 | -76 | -44 | 4.57 | 28 | -66 | -57 | 5.98 | 12 | -76 | -47 | 4.58 | 28 | -66 | -54 | 5.33 | 25 | -66 | -50 | 3.98 | | | | |
| | | | | | 15 | -74 | -40 | 4.24 | 28 | -66 | -54 | 4.32 | | | | | | | | | | | | |
| | | | | | 18 | -74 | -22 | 4.81 | 15 | -66 | -27 | 3.76 | 32 | -61 | -24 | 5.49 | 32 | -61 | -24 | 5.1 | | | | |

**Supplementary table 2 - Gestures > baseline.** Hemisphere and anatomical regions, MNI coordinates and Z score of peak activations. Voxelwise threshold $p<0.001$, clusterwise threshold $p<0.05$ FDR-corrected over the whole brain. Comparisons between groups are masked by the considered activation in the first group. L=left; R=right; a=anterior; p=posterior; LOTC=lateral occipitotemporal cortex; STS=superior temporal sulcus; STG=superior temporal gyrus; MTS=middle temporal sulcus; MTG=middle temporal gyrus; IFG=inferior frontal gyrus; SMA=supplementary motor area; IPS=intraparietal sulcus; FEF=frontal eye field

| Region | Deaf | | | | Hearing | | | | Controls | | | | Deaf > Controls | | | | Hearing > Deaf | | | | Controls > Deaf | | | |
|---|---|---|---|---|---|---|---|---|---|---|---|---|---|---|---|---|---|---|---|---|---|---|---|---|
| | x | y | z | Z | x | y | z | Z | x | y | z | Z | x | y | z | Z | x | y | z | Z | x | y | z | Z |
| L LOTC | -32 | -94 | -7 | 4.64 | -30 | -91 | -10 | 4.38 | -32 | -94 | -12 | 4.3 | -45 | -71 | 3 | 4.24 | | | | | | | | |
| R LOTC | | | | | 32 | -94 | -7 | 5.3 | | | | | | | | | | | | | | | | |
| L aSTS/STG | | | | | -55 | -4 | -10 | 5.69 | -60 | -8 | -4 | 5.78 | | | | | -60 | -6 | -4 | 6.01 | -58 | -11 | -4 | 5.73 |
| R aSTS/STG | | | | | 55 | -8 | -10 | 5.87 | 58 | -4 | -10 | 5.17 | | | | | 48 | -31 | 6 | 4.98 | 60 | -14 | -7 | 5.8 |
| L mSTS/STG | | | | | -65 | -21 | -2 | 5.56 | -60 | -28 | 0 | 5.31 | | | | | -60 | -34 | 3 | 5.66 | -62 | -34 | 3 | 5.72 |
| R mSTS/STG | | | | | 65 | -26 | 0 | 5.67 | 60 | -11 | -4 | 5.3 | | | | | 62 | -21 | -2 | 5.4 | 45 | -24 | -2 | 4.72 |
| L pSTS/STG | | | | | | | | | -62 | -41 | 3 | 5.34 | | | | | -48 | -46 | 8 | 5.73 | | | | |
| L pSTG/SMG | | | | | | | | | | | | | | | | | | | | | -62 | -44 | 18 | 4.07 |
| R pSTS/STG | | | | | | | | | 60 | -31 | 0 | 5.77 | | | | | 55 | -8 | -10 | 5.69 | 55 | -34 | 3 | 5.09 |
| L IFG (pars triangularis) | | | | | | | | | -48 | 12 | 0 | 4.51 | | | | | | | | | -35 | 29 | -2 | 6.07 |
| L IFG (pars orbitalis) | | | | | | | | | -38 | 32 | 0 | 4.71 | | | | | | | | | | | | |
| R IFG (pars triangularis) | | | | | | | | | 45 | 24 | 0 | 4.46 | | | | | | | | | 50 | 19 | -2 | 4.17 |
| L Precentral gyrus | | | | | -55 | -11 | 43 | 4.09 | | | | | | | | | -52 | -8 | 43 | 4.33 | | | | |
| L/R SMA | | | | | | | | | 2 | 4 | 60 | 4.81 | | | | | 0 | 9 | 58 | 4.78 | 0 | 4 | 63 | 5.21 |
| L Cerebellum | | | | | | | | | | | | | | | | | -45 | -66 | -27 | 4.47 | | | | |
| | | | | | | | | | -30 | -64 | -52 | 3.91 | | | | | | | | | -25 | -76 | -50 | 3.99 |
| R Cerebellum | | | | | | | | | | | | | | | | | 45 | -64 | -27 | 4.85 | 32 | -61 | -27 | 4.5 |
| | | | | | 30 | -66 | -52 | 4.14 | 30 | -64 | -50 | 4.26 | | | | | 28 | -66 | -50 | 5.97 | 28 | -66 | -50 | 6.07 |

**Supplementary table 3 - Lip-reading > Gestures.** Hemisphere and anatomical regions, MNI coordinates and Z score of peak activations. Voxelwise threshold $p<0.001$, clusterwise threshold $p<0.05$ FDR-corrected over the whole brain. Comparisons between groups are masked by the considered activation in the first group. L=left; R=right; a=anterior; m=middle; p=posterior; LOTC=lateral occipitotemporal cortex; STS=superior temporal sulcus; STG=superior temporal gyrus; SMG=supramarginal gyrus; IFG=inferior frontal gyrus; SMA=supplementary motor area

| | Deaf | | | | Hearing | | | | Controls | | | | Deaf > Controls | | | | Deaf > Hearing | | | | Controls > Deaf | | | |
|---|---|---|---|---|---|---|---|---|---|---|---|---|---|---|---|---|---|---|---|---|---|---|---|---|
| Region | x | y | z | Z | x | y | z | Z | x | y | z | Z | x | y | z | Z | x | y | z | Z | x | y | z | Z |
| L Calcarine sulcus | -10 | -101 | 0 | 4.3 | | | | | -5 | -101 | 3 | 3.77 | | | | | | | | | | | | |
| R Calcarine sulcus | 10 | -101 | 3 | 6.26 | 12 | -98 | 8 | 5.89 | | | | | | | | | | | | | | | | |
| R Middle occipital gyrus | 20 | -96 | 13 | 6.04 | | | | | | | | | | | | | | | | | | | | |
| L Superior occipital gyrus | | | | | -22 | -74 | 33 | 5.52 | | | | | | | | | | | | | | | | |
| R Superior occipital gyrus | | | | | 28 | -74 | 33 | 4.88 | 25 | -78 | 33 | 4.55 | | | | | | | | | | | | |
| L LOTC | -48 | -71 | 3 | 5.73 | -48 | -71 | 0 | 6.07 | -45 | -71 | 3 | 6.38 | | | | | | | | | -38 | -84 | -2 | 4.85 |
| | | | | | | | | | | | | | | | | | | | | | -45 | -71 | 3 | 4.24 |
| R LOTC | 48 | -66 | 3 | 6.27 | 50 | -68 | -2 | 5.79 | 52 | -66 | -2 | 6.01 | | | | | | | | | | | | |
| L LOT sulcus | -45 | -44 | -22 | 4.44 | -40 | -64 | -2 | 5.31 | | | | | | | | | | | | | | | | |
| R LOT sulcus | | | | | 42 | -54 | -12 | 3.75 | | | | | | | | | | | | | | | | |
| L Fusiform gyrus | | | | | | | | | -40 | -51 | -14 | 3.94 | | | | | | | | | | | | |
| R Fusiform gyrus | | | | | | | | | -15 | -88 | -12 | 4.96 | | | | | | | | | | | | |
| L aSTS/STG | | | | | | | | | | | | | -58 | -11 | -4 | 5.73 | -60 | -6 | -4 | 6.01 | | | | |
| R aSTS/STG | | | | | | | | | | | | | 60 | -14 | -7 | 5.8 | 60 | -11 | -10 | 5.47 | | | | |
| L mSTS/STG | | | | | | | | | | | | | -62 | -34 | 3 | 5.72 | -60 | -34 | 3 | 5.66 | | | | |
| R mSTS/STG | | | | | | | | | | | | | 45 | -24 | -2 | 4.72 | 62 | -21 | -2 | 5.4 | | | | |
| L pSTS/STG | -50 | -46 | 8 | 4.68 | | | | | | | | | | | | | -48 | -46 | 8 | 5.73 | | | | |
| | | | | | | | | | | | | | | | | | -58 | -38 | 18 | 4.03 | | | | |
| R pSTS/STG | 60 | -34 | 18 | 4.37 | 65 | -41 | 18 | 3.91 | | | | | 55 | -34 | 3 | 5.09 | 48 | -31 | 6 | 4.98 | | | | |
| L Supramarginal gyrus | -55 | -41 | 23 | 5.23 | | | | | | | | | | | | | | | | | | | | |
| R Supramarginal gyrus | | | | | 55 | -34 | 26 | 4.72 | | | | | | | | | | | | | | | | |
| L IFG (pars opercularis) | -48 | 6 | 13 | 4.13 | | | | | | | | | -48 | 12 | 10 | 4.48 | -52 | 12 | 18 | 3.99 | | | | |
| | -38 | 6 | 26 | 3.98 | | | | | | | | | | | | | | | | | | | | |
| L IFG (pars triangularis) | -35 | 29 | -2 | 3.95 | | | | | | | | | -35 | 29 | -2 | 6.07 | | | | | | | | |
| R IFG (pars orbitalis) | | | | | | | | | | | | | | | | | | | | | | | | |
| L Insula | | | | | | | | | | | | | -30 | 22 | 8 | 4.85 | -38 | 29 | -2 | 4.07 | | | | |
| R Insula | | | | | | | | | | | | | 38 | 26 | 8 | 4.92 | 38 | 26 | 8 | 4.14 | | | | |
| L Precentral gyrus | -42 | -1 | 50 | 4.43 | -52 | 2 | 33 | 5 | | | | | | | | | -52 | -8 | 43 | 4.33 | | | | |
| R Postcentral gyrus | 32 | -36 | 48 | 3.82 | | | | | 32 | -36 | 53 | 4.73 | | | | | | | | | | | | |
| L SMA | -5 | 2 | 68 | 4.52 | | | | | | | | | | | | | | | | | | | | |
| L/R SMA | | | | | | | | | | | | | 0 | 4 | 63 | 5.21 | 0 | 9 | 58 | 4.78 | | | | |
| R Middle frontal gyrus | 40 | -4 | 56 | 4.25 | | | | | | | | | | | | | | | | | | | | |
| L IPS | | | | | -35 | -41 | 46 | 6.32 | -18 | -58 | 58 | 5.7 | | | | | | | | | | | | |
| | | | | | -22 | -64 | 60 | 7.06 | -25 | -58 | 63 | 4.46 | | | | | | | | | | | | |
| R IPS | | | | | 30 | -48 | 56 | 5.32 | | | | | | | | | | | | | | | | |
| L FEF | | | | | -25 | -11 | 53 | 5.67 | | | | | | | | | | | | | | | | |
| R FEF | | | | | 40 | -6 | 50 | 4.71 | | | | | | | | | | | | | | | | |
| L Cerebellum | | | | | | | | | | | | | -28 | -66 | -57 | 3.74 | | | | | | | | |
| | | | | | | | | | | | | | | | | | -45 | -66 | -27 | 4.47 | | | | |
| R Cerebellum | 28 | -66 | -50 | 4.72 | | | | | | | | | 28 | -66 | -50 | 6.07 | 28 | -66 | -50 | 5.97 | | | | |
| | 45 | -58 | -30 | 4.91 | | | | | | | | | 45 | -64 | -27 | 4.77 | 48 | -61 | -30 | 5.16 | | | | |
| | 32 | -61 | -27 | 3.96 | | | | | | | | | 32 | -61 | -27 | 4.5 | | | | | | | | |

**Supplementary table 4 - Gestures > Lip-reading.** Hemisphere and anatomical regions, MNI coordinates and Z score of peak activations. Voxelwise threshold $p<0.001$, clusterwise threshold $p<0.05$ FDR-corrected over the whole brain. Comparisons between groups are masked by the considered activation in the first group. L=left; R=right; a=anterior; m=middle; p=posterior; LOTC=lateral occipitotemporal cortex; STS=superior temporal sulcus; STG=superior temporal gyrus; IFG=inferior frontal gyrus; SMA=supplementary motor area; IPS=intraparietal sulcus; FEF=frontal eye field

| Region | Conjunction Controls & Hearing x | y | z | Z |
|---|---|---|---|---|
| L STS/STG and MTG | -55 | -18 | 6 | > 8 |
| R STS/STG and MTG | 52 | -14 | 6 | 7.74 |
| L Cerebellum | -18 | -68 | -54 | 5.11 |
| R Cerebellum | 15 | -71 | -54 | 4.69 |

**Supplementary table 5 - Audible > Silent Sentences.** Hemisphere and anatomical regions, MNI coordinates and Z score of peak activations. Voxelwise threshold p<0.001, clusterwise threshold p<0.05 FDR-corrected over the whole brain. L=left; R=right; STS=superior temporal sulcus; STG=superior temporal gyrus; MTG=middle temporal gyrus

| | Deaf | | | | Hearing | | | |
|---|---|---|---|---|---|---|---|---|
| Region | x | y | z | Z | x | y | z | Z |
| L Lingual gyrus | -8 | -96 | -10 | 4.26 | | | | |
| L IFG | | | | | -40 | 22 | 3 | 4.75 |
| L Insula | | | | | -30 | 32 | 3 | 4.27 |
| R Insula | | | | | 32 | 29 | 6 | 4.01 |

**Supplementary table 6 - Silent meaningful > Pseudo-Sentences.** Hemisphere and anatomical regions, MNI coordinates and Z score of peak activations. Voxelwise threshold p<0.001, clusterwise threshold p<0.05 FDR-corrected over the whole brain. Other groups and comparisons were non-significant. L=left; R=right; IFG=inferior frontal gyrus

(A)

| Region | x | y | z | Z |
|---|---|---|---|---|
| **Words > other categories** | | | | |
| L Calcarine sulcus | -10 | -91 | -2 | 4.72 |
| R Calcarine sulcus | 15 | -86 | -2 | 4.44 |
| L lateral occipitotemporal sulcus | -50 | -58 | -17 | 3.88 |
| L pSTS/STG | -60 | -31 | -2 | 3.94 |
| L IFG (pars opercularis) | -40 | 6 | 26 | 3.96 |
| **Faces > other categories** | | | | |
| L/R Calcarine sulcus | 2 | -84 | -10 | 4.80 |
| R Cuneus | 10 | -96 | 8 | 4.59 |
| L Fusiform gyrus | -42 | -56 | -22 | 4.97 |
| R Fusiform gyrus | 42 | -51 | -20 | 4.97 |
| R pSTS/STG | 52 | -44 | 13 | 4.08 |
| **Bodies > other categories** | | | | |
| L LOTC | -48 | -78 | 6 | 5.45 |
| R LOTC | 50 | -71 | 6 | 6.93 |

(B)

| Region | x | y | z | Z |
|---|---|---|---|---|
| **Tools > other categories** | | | | |
| R Middle occipital gyrus | 35 | -81 | 6 | 7.01 |
| R Superior occipital gyrus | 28 | -76 | 33 | 4.63 |
| L LOTC | -30 | -88 | 13 | 6.8 |
| R LOTC | 48 | -64 | -4 | 5.7 |
| L Collateral sulcus | -22 | -68 | -12 | 6.49 |
| R Collateral sulcus | 25 | -64 | -10 | 5.51 |
| | 30 | -68 | -14 | 5.74 |
| L IPS | -25 | -56 | 58 | 4.07 |
| R IPS | 30 | -56 | 63 | 4.71 |
| **Houses > other categories** | | | | |
| L Occipital pole | -25 | -86 | 20 | 6.72 |
| R Occipital pole | 15 | -96 | 3 | 7.12 |
| L Collateral sulcus | -22 | -76 | -12 | 7.16 |
| R Collateral sulcus | 25 | -78 | -7 | 6.17 |
| L Lingual gyrus | -22 | -81 | -17 | 7.01 |
| R Lingual gyrus | 18 | -84 | -12 | 7.34 |
| R Precuneus | 20 | -56 | 10 | 4.95 |

**Supplementary table 7 - Category-specific activations for the conjunction of all three groups.** Hemisphere and anatomical regions, MNI coordinates and Z score of peak activations for (A) Words, Faces and Bodies and (B) Tools and Houses. Voxelwise threshold $p<0.001$, clusterwise threshold $p<0.05$ FDR-corrected over the whole brain. No comparison showed significant activation. L=left; R=right; p=posterior; STS=superior temporal sulcus; STG=superior temporal gyrus; IFG=inferior frontal gyrus; LOTC=lateral occipitotemporal cortex; IPS=intraparietal sulcus

| | Region | | | | | | | | | | | | | | | |
|---|---|---|---|---|---|---|---|---|---|---|---|---|---|---|---|---|
| | **L Collateral sulcus** | | | | **L pSTS/STG** | | | | **R pSTS/STG** | | | | **R mSTS/STG** | | | |
| | **x** | **y** | **z** | **X** | **x** | **y** | **z** | **X** | **x** | **y** | **z** | **X** | **x** | **y** | **z** | **X** |
| Words > baseline | | | | | | | | | 65 | -36 | 8 | 6.12 | 65 | -16 | -2 | 6.11 |
| Faces > baseline | -22 | -68 | -2 | 4.43 | -58 | -44 | 10 | 4.48 | 62 | -36 | 8 | 6.53 | 65 | -16 | -2 | 5.44 |
| Bodies > baseline | -22 | -68 | -2 | 4.37 | | | | | 62 | -34 | 8 | 6.08 | | | | |
| Tools > baseline | | | | | | | | | 68 | -36 | 8 | 6.07 | | | | |

**Supplementary table 8 – Comparison Deaf > Hearing + Controls (masked by Deaf) for each category > baseline.** Hemisphere and anatomical regions, MNI coordinates and Z score of peak activations. Voxelwise threshold p<0.001, clusterwise threshold p<0.05 FDR-corrected over the whole brain. No Houses > baseline comparison showed significant activation. L=left; R=right; p=posterior; m=middle; STS=superior temporal sulcus; STG=superior temporal gyrus
